# Short-term diet intervention alters the small non-coding RNA (sncRNA) landscape of human sperm

**DOI:** 10.1101/2021.07.08.451257

**Authors:** Candida Vaz, Alexandra J Kermack, Mark Burton, Pei Fang Tan, Jason Huan, Tessa P X Yoo, Kerry Donnelly, Susan J Wellstead, Helena L Fisk, Franchesca D Houghton, Sheena Lewis, Yap Seng Chong, Peter D Gluckman, Ying Cheong, Nicholas S Macklon, Philip C Calder, Anindya Dutta, Keith M Godfrey, Pankaj Kumar, Karen A Lillycrop, Neerja Karnani

**Author notes:** represent joint authorships (Candida Vaz, Alexandra J Kermack, Mark Burton, Keith M Godfrey, Pankaj Kumar, Karen A Lillycrop, Neerja Karnani).

## Abstract

Offspring health outcomes are often linked with epigenetic alterations triggered by maternal nutrition and intrauterine environment. Strong experimental data also link paternal preconception nutrition with pathophysiology in the offspring, but the mechanism(s) routing the effects of paternal exposures remain elusive. Animal experimental models have highlighted small non-coding RNAs (sncRNAs) as potential regulators of paternal effects, though less is known about the existence of similar mechanisms in human sperm. Here, we first characterised the baseline sncRNA landscape of human sperm, and then studied the effects of a 6-week diet intervention on their expression profile. Baseline profiling identified 5’tRFs, miRNAs and piRNAs to be the most abundant sncRNA subtypes, primarily expressed from regulatory elements like UTRs, CpG-rich regions and promoters. Expression of a subset of these sncRNAs varied with age, BMI and sperm quality of the donor. Diet intervention enriched in vitamin D and omega-3 fatty acids showed a marked increase of these nutrients in circulation and altered the sperm sncRNA expression. These included 3 tRFs, 15 miRNAs and 112 piRNAs, with gene targets involved in fatty acid metabolism, vitamin D response (LXR/RXR activation, TGF-beta and Wnt signaling), and transposable elements. These findings provide evidence that human sperms are sensitive to alterations in exposures such as diet, and sncRNAs capture the epigenetic imprint of this change. Hence changes to paternal nutrition during preconception may improve sperm quality and offspring health outcomes. To benefit future research, we developed iDad_DB, an open access database of baseline and diet-altered sncRNA in human male germline.

## Introduction

Inheritance of genetic material has long been acknowledged as the primary route of transmitting health related information from parents to the offspring. However, emerging evidence from the DOHaD (Developmental Origins of Health and Disease) field identifies epigenetic mechanisms as an additional route of transgenerational inheritance, especially in transferring the effects of parent’s lifestyle and environmental exposures, with implications for the offspring’s risk of non-communicable diseases (NCDs) in later life. Epidemiological, clinical and basic science research has characterised the period around conception as being critical in the processes linking parental environmental exposures with the health of the next generation (Fleming et al. 2018). Since these exposures and the related epigenetic perturbations constitute modifiable factors, they can serve as promising targets for developing effective interventions against NCDs. Current DOHaD research is primarily focused at understanding the role of maternal factors, such as nutrition, in shaping the development of the offspring, but less is known about the mechanisms involved in the transmission of paternal exposures through the male germline.

A variety of lifestyle and environmental factors ranging from nutrition (Carone et al. 2010; Ng et al. 2010; Ng et al. 2014), stress (Crews et al. 2012; Skinner 2014; McCreary et al. 2016) and exposure to toxins such as fungicides, pesticides (Manikkam et al. 2012; Manikkam et al. 2013; Skinner et al. 2013; Skinner et al. 2018) and air pollution (Deng et al. 2016) are known to alter semen concentration and sperm quality. It is also becoming apparent that these exposures may leave an imprint on the sperm epigenome, that may be transmitted to the new life at conception (Donkin and Barres 2018; Siddeek et al. 2018; Bedi et al. 2019). Studies from rodent models have provided some insights into the mechanisms involved in mediating these effects to the offspring (Fullston et al. 2013; Gapp et al. 2014; Fleming et al. 2018). These primarily involve alterations to DNA methylation and small non-coding RNA (sncRNA) expression (Siddeek et al. 2018). Sperm sncRNA expression in rodents is known to be sensitive to paternal exposures such as diet, and capable of signaling it’s effects to the offspring (Donkin and Barres 2018).

sncRNAs include a variety of small RNA species, that are typically <100 nucleotides long, and regulate diverse cellular processes. Their size and structure can give sncRNAs greater stability, and mobility across tissues, making them ideal candidates for intercellular signaling and gene regulation. These characteristics also make sncRNAs pivotal regulators in developmental and physiological processes. MicroRNAs (miRNAs) are an example of such small RNA species, that are typically 22-24 nucleotides long, and can post-transcriptionally regulate gene expression by base-pairing with complementary sequences on mRNA. They can modulate other epigenetic regulators, and conversely themselves be targets of epigenetic regulation (Shukla et al. 2011; Gruber and Zavolan 2013). Studies in mouse models have shown that stress (Rodgers et al. 2015) and diet (Grandjean et al. 2015) can alter miRNA expression in the male germline. Furthermore, injecting the environment responsive sperm miRNAs into the zygote/embryo can remodel offspring gene expression, and propagate paternal phenotypes to the subsequent generation.

tRNA-derived small RNAs (tsRNAs) (Peng et al. 2012), also known as tRNA-derived fragments (tRFs) (Cole et al. 2009; Lee et al. 2009; Haussecker et al. 2010), constitute another class of sncRNAs implicated in the trans-generational epigenetic inheritance in rodent model systems. One of the earliest discovered classes of tRFs were the tiRNA (tiR), which are typically 31–40 nucleotides long. The other tRF classes constitute the relatively shorter subtypes, that are 14–30 nucleotides long and map to the 5’ and 3’ ends of the mature or primary tRNA transcripts. tRF-5 and tRF-3 are generated from 5′ and 3′ ends of the mature tRNAs, while tRF-1 is generated from the 3′ end of the primary tRNA transcripts. Depending upon their length, tRF-5 and tRF-3 can be further divided into a, b or c sub-classes (Kumar et al. 2015; Kumar et al. 2016). This heterogeneity in the tRNA derivatives provides a sense of complexity in gene regulation mediated by these small RNA molecules.

Chen *et al*., (Chen et al. 2016) using a rodent model system, reported that feeding a high fat diet (HFD; 60% fat) to male mice altered the expression of their sperm tsRNA. Injection of these sperm-derived tsRNAs from the HFD-fed males into normal zygotes generated metabolic disorders in the F1 offspring. Similarly, Sharma *et al*., (Sharma et al. 2016) also showed the effects of diet (low protein) on the levels of miRNAs and tsRNA expression in mature mouse sperms. In a recent study by Natt *et al*., (Natt et al. 2019) the short-term effects of a 2-step diet intervention on human sperm was demonstrated. Their study reported that human sperm is highly sensitive to a sugar-rich diet, which affected sperm motility and biogenesis of tsRNAs, suggesting an interplay between nutrition and male reproductive health.

In addition to miRNA and tRFs, piRNAs also constitute an abundant class of sncRNA in the male germline. piRNAs range from 25-33 nucleotides in size, with 5’ sequences enriched in uridine and 2’O-methyl modifications at the 3’end. The primary function of piRNA is to silence the transposable elements to protect genome integrity (Czech and Hannon 2016; Ernst et al. 2017), but they have also been reported to play a role in DNA methylation and mRNA stability. piRNA can also respond to various environmental stressors, raising the possibility of existing mechanisms that directly signal the effects of environmental changes to piRNA biogenesis machinery (Belicard et al. 2018). However, there are currently no studies reporting the effects of diet on piRNA expression in human sperm.

As is evident from animal model systems, sperm borne sncRNAs may provide an epigenetic route for inter-generational signaling of paternal exposures. However, little is known about the existence of similar or related mechanisms in humans. In this study, we aimed to characterise the baseline sncRNA landscape of human sperm, and study the effects of a paternal diet, enriched in vitamin D and omega-3 fatty acids on sncRNA expression.

## Results

Sperm sncRNA profiling was performed on a subset (N=17) of participants enrolled in the PREconception dietary suPplements in Assisted REproductionin (PREPARE) study in the UK (N=111 recruited and 102 completed the study; see Methods for details) (Kermack et al. 2014). Sperm samples were collected twice during the study, i.e. before and 6 weeks after the dietary intervention (visits MV1 and MV4, **Supplemental File S1**). Of the 17 participants selected for sncRNA analysis, these were the only samples with sufficient RNA both pre and post intervention for small RNAseq analysis; 9 were participants in the control arm, and 8 from the intervention arm of the trial. The intervention group received an olive oil based diet plus a supplemented drink enriched in vitamin D and omega-3 fatty acids for 6 weeks, while the control group received a sunflower oil based diet and a placebo drink for the same duration of time (**Figure 1A)**. Small RNA sequencing was performed on the total RNA extracted from the sperm samples. Number of sequence reads per sample ranged from 10M to 21M, and the sequence length ranged from 10 – 46 nucleotides. sncRNAseq reads were mapped to the human genome and annotated into different subtypes using the Genboree software and additional miRNA and tRF annotators detailed in the methods.

**Figure 1.**
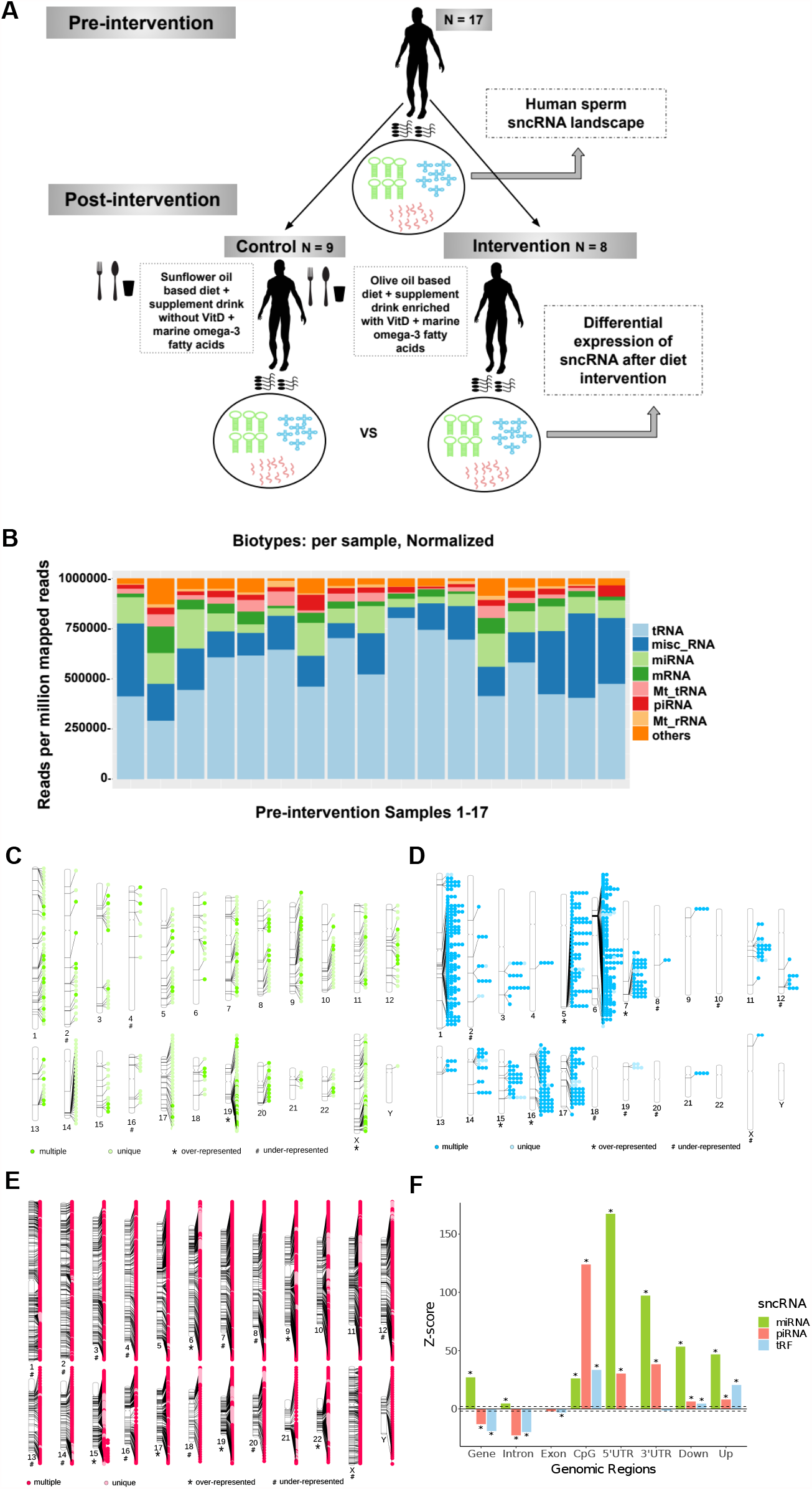
Study design and baseline characterization of sncRNA landscape of human sperms. (A) Illustration of the study design indicating the number of subjects and sncRNA analysis conducted in the pre and post-intervention groups. (B) Relative abundance of sncRNA subtypes identified during the baseline characterization of sncRNA landscape of sperm samples collected at the pre-intervention visit. tRFs were the most abundant sncRNA, followed by miRNA and piRNA. Small amounts of mitochondrial tRNAs and rRNAs were also present. The misc_RNA included additional small RNA subtypes and fragmented mRNA. ‘Other’ includes small RNAs identified in this study, but not yet annotated in available sncRNA databases. (C) Phenogram showing chromosome based enrichment analysis of 401 baseline miRNAs. Lighter shade of green represents the miRNA expressed from unique genomic locations, and the darker shade represents the miRNA that mapped to multiple sites in the genome. ^*^: chromosomes with over-represented miRNA expression (Z-score ≥ 2). #: chromosomes with under-represented miRNA expression (Z-score ≤ -2). (D) Phenogram showing chromosome-based enrichment analysis of 143 baseline tRFs. Lighter shade of blue represents the tRF expressed from unique genomic locations, and the darker shade represents the tRF that mapped to multiple sites in the genome. ^*^: chromosomes with over-represented tRF expression (Z-score ≥ 2). #: chromosomes with under-represented tRF expression (Z-score ≤ -2). (E) Phenogram showing chromosome-based enrichment analysis of 2290 baseline piRNAs. Lighter shade of pink represents the piRNA expressed from unique genomic locations, and the darker shade represents the piRNA that mapped to multiple sites in the genome. ^*^: chromosomes with over-represented piRNA expression (Z-score ≥ 2). #: chromosomes with under-represented piRNA expression (Z-score ≤ -2) (F) Genomic annotation enrichment of the sperm baseline sncRNAs. The genomic regions analysed comprised of genes, introns exons, CpG regions, 5’UTRs, 3’UTRs, 2Kb downstream of transcription termination site (Down) and 2Kb upstream of transcription start site (Up).

### Pre-intervention sncRNA landscape of human sperm

sncRNA profiles of the 17 pre-intervention sperm samples were used to determine the baseline sncRNA landscape of the subjects in the study. tRFs were the most abundant sncRNA species detected in sperm, followed by miRNA and piRNAs. Small amounts of mitochondrial tRNA and rRNA were also detected (**Figure 1B**).

High stringency analysis, based on the criterion of detecting sncRNA expression in all 17 subjects, identified a total of 143 tRFs, 401 miRNAs, and 2290 piRNAs (**Supplemental File S2**). Most of the sperm tRFs (63.6%, 91/143) originated from multiple genomic loci, while miRNA (86.3%, 346/401) and piRNA (79.7%, 1826/2290) were predominantly expressed from unique genomic locations (**Figures 1C-E**).

Chromosome based enrichment analysis revealed expression of a higher number of miRNA from chromosomes 19 and X (**Figure 1C** and **Supplemental Figure S1**). Likewise, a higher number of tRF expression was observed from 4 chromosomes (i.e. 5, 7, 15 and 16; **Figure 1D** and **Supplemental Figure S2**), and piRNA from 6 chromosomes (6, 9 15, 17, 19 and 22; **Figure 1E** and **Supplemental Figure S3**). We also noted expression of miRNA and piRNA from the Y chromosome. Genomic annotation analysis revealed enrichment of the sperm sncRNAs expression from the regulatory regions of the genome, and depletion from the coding exons. Majority of the miRNAs originated from the UTRs, while tRFs and piRNA were predominantly expressed from CpG rich regions of the sperm genome (**Figure 1F**). There was also enrichment of sncRNAs expression from the promoters (2Kb upstream of the transcription start site) and 2Kb downstream of the transcription termination sites. Introns showed a slight enrichment of miRNA expression, but were generally depleted of tRFs and piRNA expression.

Ninety-seven percent of the tRFs were derived from the 5′-end of tRNA, including 5′-tiRs and 5′-tRFs (**Supplemental File S2**). In contrast, 3′-tiRs, 3′-tRFs, and 1’-tRFs were underrepresented in sperm. For each tRF id, the most abundant genomic sequence (having the highest read count) was extracted and compared across the pre-intervention samples, demonstrating sequence as well as length conservation across the samples. Around 80% of the most abundant tRF sequences were almost the same length (+-2 nucleotides) across the samples. The most abundant sequence was taken as the representative sequence for a tRF. (**Supplemental File S3**).

422/2290 (18.4%) baseline piRNAs were located within the transposable elements (LINE:106, SINE:66, LTR:195, Satellites:3, Others:53), and 17/2290 (0.7%) originated from tRNAs (**Supplemental File S4**). Transposable elements (TE) associated with piRNA comprised of LINE elements such as L1, L2 and L3, SINE elements such as the Alu family, LTRs such as ERV1, ERVL, ERVL-MaLR, Gypsy, DNA transposons such as Tigger3b|DNA|TcMar-Tigger and MER|DNA|hAT-Charlie, and Satellite elements (**Supplemental File S5)**. 1041 of the remaining baseline piRNAs mapped to the genes, long intergenic non-coding RNA (lincRNA) and pseudogenes (**Supplemental File S5)**. We found 234 genes related to the 1041 baseline piRNAs to be enriched in pathways such as Golgi and Endomembrane system organization (**Supplemental Table S1**)

### Factors contributing to inter-individual variation in the sncRNA profiles

To investigate the factors contributing to the inter-individual variation in sperm sncRNA profiles of the subjects in the pre-intervention group, we studied the effects of age, BMI, and sperm fertility measures such as sperm concentration and sperm motility (**Supplemental File S1**). miRNA expression was found to be significantly associated with age and sperm concentration (FDR corrected p-value <0.05, **Supplemental File S6**). Expression levels of 56 miRNAs (15 up-regulated and 41 down-regulated) were altered with age (**Figure 2A(i)**), and 127 miRNAs (57 up-regulated and 70 down-regulated) with sperm concentration (**Figure 2A(ii))**. To ascertain the genes targeted by miRNAs associated with age and sperm concentration, experimentally validated and/or high scoring predicted targets of these miRNAs were identified. Pathway enrichment analysis of the target genes by IPA core analysis (details under Methods) identified the age dependent miRNA targets to be enriched in the pathways known to be linked with ageing, such as mTOR signaling, apoptosis and p53 signaling, telomerase signaling, PI3K/AKT signaling, Wnt/B-catenin and TGF-B signaling (**Supplemental Figure S4**). Likewise, the gene targets for sperm concentration associated miRNAs were enriched in multiple cellular pathways including those linked with male fertility and spermatogenesis, such as androgen signaling, sertoli and germ cell junction signaling, integrin and Protein Kinase A signaling (**Supplemental Figure S5)**. Sperm tRF expression was significantly associated with BMI and sperm concentration (FDR corrected p-value <0.05). Six tRFs were downregulated in males with high BMI (**Figure 2B(i)** and **Supplemental Table S2A**), and another six tRFs (3 up-regulated and 3 down-regulated) showed altered expression in association with sperm concentration (**Figure 2B(ii)** and **Supplemental Table S2B**). All 12 tRFs were 5’-end derivatives of their respective primary tRNA. piRNA expression was associated only with sperm concentration, with 1 piRNA up-regulated and 2 down-regulated (**Figure 2C** and **Supplemental Table S2C**).

**Figure 2.**
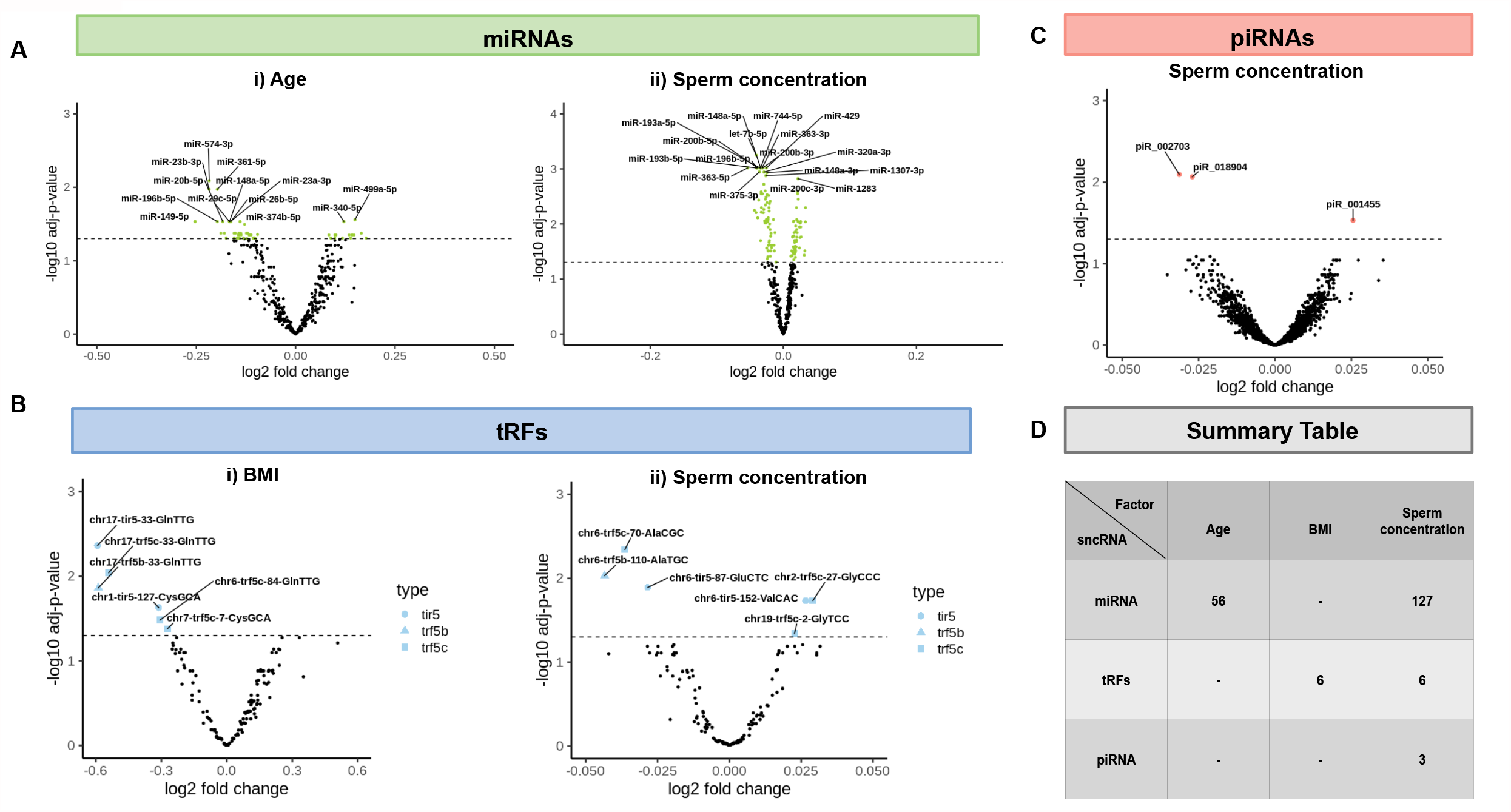
sncRNA associated with age, BMI and sperm concentration. (A) Volcano plots showing miRNAs associated with (i) age and (ii) sperm concentration. Y-axis represents the –log10 adjusted p-value of association between the miRNA and the tested paternal factor, while X-axis represents the log2 fold change in the expression of miRNA per unit change in age, or sperm concentration. Dashed horizontal line represents the significance threshold (FDR corrected -log10 p-value = 1.3) for association analysis. Green dots above this threshold represent the miRNAs whose expression were significantly altered with age and sperm concentration. Top miRNA from both the association studies are labeled in their respective volcano plots. (B) Volcano plots showing tRFs associated with (i) BMI and (ii) sperm concentration. Dashed horizontal line represents the significance threshold (-log10 = 1.3) for the association analysis, and the blue colored shapes represent the different tRF subtypes whose expression were significantly altered in association with BMI and sperm concentration. (C) Volcano plots showing piRNAs (pink dots) associated with sperm concentration. (D) Table summarizing the number of sncRNAs subtypes associated with age, BMI and sperm concentration.

In summary, sperm concentration had more generalised impact on the different sncRNA subtypes, while age and BMI were associated with changes in miRNA and tRF expression, respectively (**Figure 2D**). This pre-intervention assessment of inter-individual variation in sncRNA profiles also helped identify the factors that could confound the post-intervention analysis. Sperm motility was not found to be a significant factor in our study, but since it has been previously reported to affect sncRNA expression (Capra et al. 2017), we included it as an additional covariate in the subsequent analysis.

### Effects of diet intervention on vitamin D and omega-3 fatty acid levels in circulation

To study the effects of diet intervention, we measured the levels of vitamin D and omega-3 fatty acids in blood samples collected before and 6 weeks post-intervention. To also assess the sampling bias, blood measures of 17 subjects included in sncRNA analysis were compared with the full dataset (N=102) from the PREPARE trial.

In the full dataset, there was a significant increase (p-value 1.9e^-13^; **Figure 3A(i)**) in the serum vitamin D concentrations of the participants in the intervention group (mean change = 83 nmol/l) as compared to the control group (mean change = 2 nmol/l). sncRNA sample subset also reflected the same trend (intervention group mean change = 109 nmol/l, control group mean change = 8 nmol/l; p-value 9.9e^-05^, **Figure 3A(ii)**). Likewise, diet intervention also showed a significant increase in the % of EPA (full set p-value < 2.2e^-16^ and sncRNA subset p-value 2.7e^-07^, **Figures 3B (i & ii)**) and % DHA (full set p-value < 2.2e^-16^ and sncRNA subset p-value 9.6e^-06^; **and Figures 3C (i & ii)**) in RBCs in both the full and sncRNA subset. Increase in the omega-3 fatty acids and vitamin D levels in the circulation provides evidence for the effectiveness of the 6-week dietary intervention. We also tested if the intervention altered the EPA and DHA composition of the seminal plasma and sperm. Percentage of EPA measured in the sperm (0.08 % (±1.57) vs 0.06 % (±1.57), p=0.007) and seminal plasma (0.12 % (±1.82) vs. 0.09 % (±2.05), p=0.032) was higher in the treatment compared to the control group, however no difference was seen in the percentage of DHA in sperm (15.43 % (±2.64) vs. 13.85 % (±1.89), p=0.379) or seminal plasma (8.99 % (±3.84) vs. 9.24 % (±4.45), p=0.764). Since the physiological changes in fatty acid levels need not always be reflected in the lipid composition of semen and sperm, effects of altered paternal nutrition might still be captured and signaled to the male germline via an epigenetic route.

**Figure 3.**
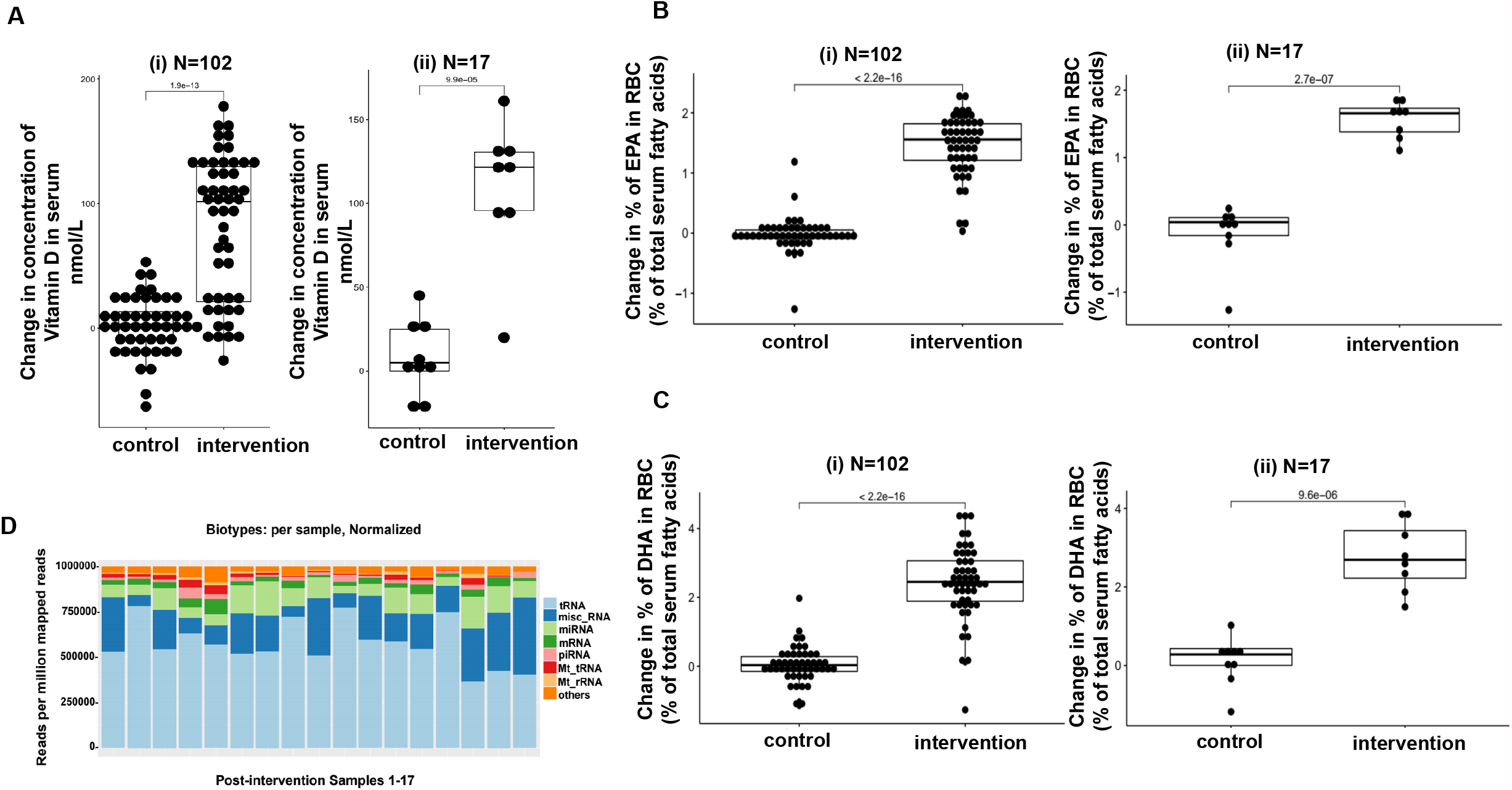
Effects of diet intervention on vitamin D and omega-3 fatty acid levels in circulation. (A) Boxplots comparing the change (pre vs post-intervention) in serum vitamin D concentration of subjects in the control and intervention groups (nmol/l). Data plots for both (i) full study (N=102) and the (ii) sncRNA subset (N=17). (B) Boxplots comparing the change (pre vs post-intervention) in the % EPA composition of RBCs of subjects in the control and intervention groups. Data plots for both (i) full study and the (ii) sncRNA subset. (C) Boxplots comparing the change (pre vs post-intervention) in the % DHA composition of RBCs of subjects in the control and intervention groups. Data plots for both (i) full study and the (ii) sncRNA subset. (D) Relative abundance of sncRNA subtypes post-intervention.

### Effect of dietary intervention on sperm sncRNA expression

We next studied the effects of dietary intervention on the sncRNA profile. Similar to the pre-intervention sncRNA composition of sperm, the post-intervention sncRNA profiles also showed a high abundance of tRFs, followed by miRNAs and piRNAs (**Figure 3D**), indicating no global changes to the relative abundance of small RNA subtypes. However, we did identify differentially expressed sncRNAs between the control and intervention groups. **Supplemental File S7** provides the list of these differentially expressed sncRNAs detected after adjusting for the effects of age, BMI, sperm concentration and motility.

#### Differentially expressed miRNAs (DE-miRNA)

mirDeep2 was used to characterise sperm miRNA expression profiles of the subjects in the control and intervention groups and identify the DE-miRNAs. To study the specific effects of diet, we adjusted the analysis for the confounding effects of age, BMI, sperm motility and sperm concentration. Fifteen miRNAs were identified to be differentially expressed between the control and intervention groups, with log2(1.5) fold change and –log10 p-value ≥ 2 (**Figures 4A & B, Supplemental Table S3A** and **Supplemental File S7**). DIANA miRPath identified 7 of these 15 miRNAs to target genes in fatty acid biosynthesis and metabolism pathways (**Figure 4C**). For example, hsa-miR-506-3p had predicted target sites in 8 genes namely: *ACAA2, ACSL1, CPT1A, ELOVL5, HADH, OXSM, PECR* and *SCD*, followed by hsa-miR-513a-3p and hsa-miR-513c-3p targeting 5 genes: *ACADSB, ACOX1, ELOVL5, FASN, PTPLB* (**Figure 4C**). Additional IPA analysis of the genes targeted by 7/15 DE-miRNA reconfirmed the enrichment of specific pathways related to fatty acid metabolism, such as fatty acid β-oxidation, mitochondrial – L carnitine shuttle pathway, stearate biosynthesis, fatty acid activation, γ-linolenate biosynthesis, palmitate biosynthesis, fatty acid biosynthesis initiation and oleate biosynthesis (**Figure 4D**). Besides fatty acid metabolism pathways, the LXR/RXR activation, TGF-beta signaling, Hippo signaling and Wnt signaling pathways were also found to be significantly enriched (**Supplemental Table S4**).

**Figure 4.**
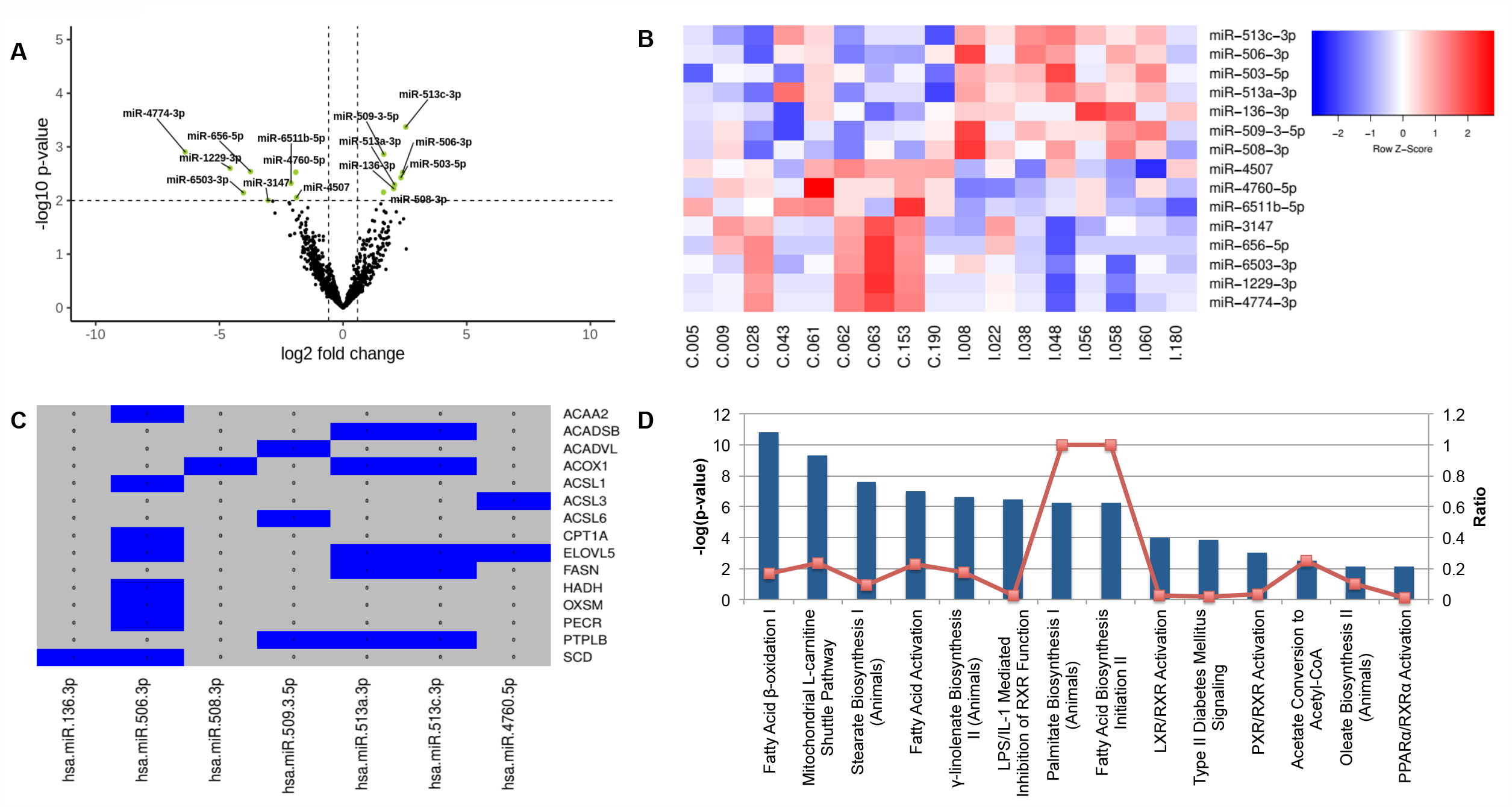
Effect of diet intervention on miRNA expression. (A) Volcano plot showing differentially expressed miRNAs (intervention vs control) identified after adjusting for age, BMI, sperm concentration and motility. Horizontal dashed line represents the -log10 p-value = 2 significance threshold of association, and the vertical dashed lines represent the log2(1.5) fold change. (B) Heatmap of miRNAs differentially expressed between the control and the intervention groups. Each row represents the expression z score for the differentially expressed miRNA ID indicated on the right, color changes from white to red indicate the magnitude of increase in expression (upregulation), while changes from white to blue indicate the magnitude of downregulation. Each column represents the de-identified subject IDs where those initiating with ‘C’ refer to the subjects in the control group and ‘I’ refers to the subjects in the intervention group. (C) Differentially expressed miRNAs and their corresponding target genes predicted by DIANA miRPath. Blue color (1) indicates the presence of the miRNA-target interaction, while grey color (0) indicates the absence of a predicted interaction. (D) Ingenuity Pathway Analysis (IPA) of genes targeted by differentially expressed miRNA. Pathways with the most over represented genes and a enrichment -log10 p-value > 2 are plotted by the decreasing order of significance. The dotted red line represents the ratio of the miRNA target genes to the total number of genes known to be present in that pathway.

#### Differentially expressed tRFs (DE-tRFs)

Expression profiling of tRNA derived fragments (tRFs) was done using the analysis pipeline developed by Kumar *et al*., (Kumar et al. 2015). Quantitative expression of 8 different categories of tRNA derived tRFs (5’tir, tRF-5 (a/b/c), tRF-3(a/b/c) and tRNA trailers (tRF-1)) was performed. After adjusting for age, BMI, sperm motility and sperm concentration, 3 tRFs were found to be differentially expressed between the intervention and control groups (log2(1.5) fold change and –log10 p-value ≥ 2; **Figure 5A & B, Supplemental File S7** and **Supplemental Table S3B)**. These three differentially expressed tRFs included derived trf5b-2-TyrGTA, tir5-29-CysGCA and trf5b-24-AlaAGC.

**Figure 5.**
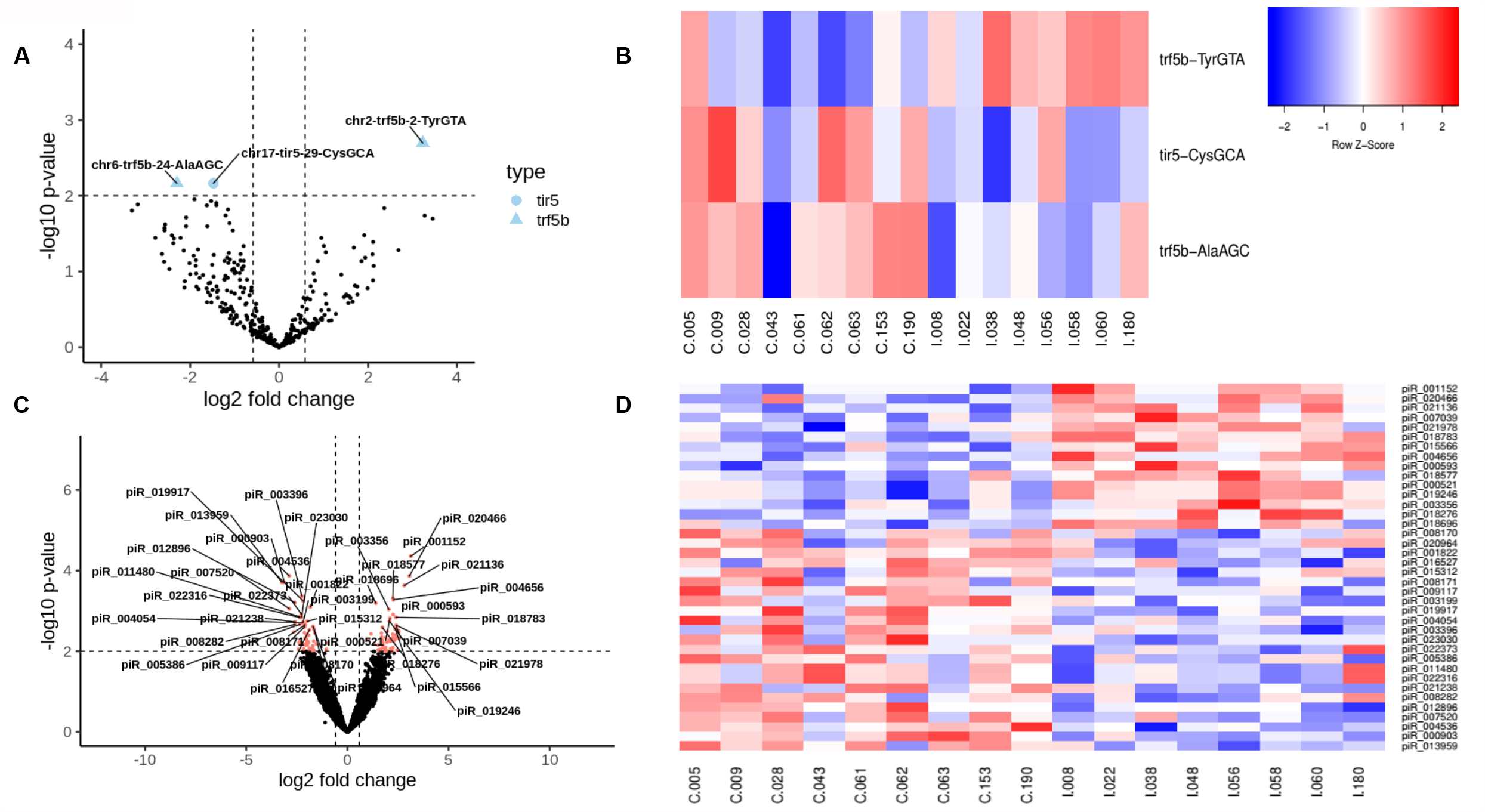
Effect of diet intervention on tRF and piRNA expression. (A) Volcano plot showing differentially expressed tRFs (intervention vs. control) identified after adjusting for age, BMI, sperm concentration and motility. Horizontal dashed line represents the -log10 p-value = 2 significance threshold of association, and the vertical dashed lines represent the log2(1.5) fold change. (B) Heatmap of tRFs differentially expressed between the control and intervention groups. (C) Volcano plot showing the differentially expressed piRNAs (intervention vs. control). (D) Heatmap of differentially expressed piRNA. Due to space restriction this heatmap is provided only for the top most significant piRNA labeled in figure 5(C).

#### Differentially expressed piRNAs (DE-piRNAs)

Differential expression analysis of piRNA adjusted for age, BMI, sperm motility and sperm concentration identified 112 piRNAs (log2(1.5) fold change and –log10 p-value ≥ 2) to differ in expression between the intervention and control groups (**Figure 5C & D** and **Supplemental File S7**). 10.7% (12/112) of these DE-piRNAs were located within the transposable elements (LINE:4, SINE:2, LTR:4, Others:2), and another 48.2% (54/112) overlapped with genomic regions encoding genes or LINC RNA (**Supplemental Files S8** and **S9**).

### iDad_DB, an open source database of sperm sncRNAs

For easy access, visualisation and future use of the sperm sncRNAs identified in this study, we developed an open access database named iDad_DB (https://idaddb.sics.karnanilab.com/). This database contains baseline, paternal factors and diet intervention responsive sncRNA identified in this study.

## Discussion

The sperm epigenome is sensitive to paternal lifestyle and environment, and can potentially transfer its beneficial or detrimental effects to the future offspring. This study provides a high resolution mapping of three most abundant sncRNA species (miRNAs, tRFs and piRNAs) in human sperm, and the effects of a healthy short-term dietary intervention on their expression. In the intervention arm of the study the participants were given a diet enriched in vitamin D and omega-3 fatty acids, known for their anti-inflammatory and beneficial roles in metabolism (Calder 2018). Despite the relatively short 6-week duration of the intervention, there was a significant increase in the levels of vitamin D, EPA and DHA in the circulation, and a change in the sperm sncRNA expression profile. This study also generated a comprehensive baseline sncRNA landscape of human sperm and identified the small RNAs associated with age, BMI and sperm quality. Finally, we developed an open access database, iDad_DB, for easy access and future use of sperm sncRNA profiles generated in the study.

Baseline characterisation of the sncRNA landscape in human sperm revealed multiple subtypes of small RNA, with varying abundance. These sncRNA were predominantly expressed from the regulatory regions of the genome, such as the UTRs. CpG rich sites and promoters suggesting their potential role in chromatin remodeling and transcriptional regulation in sperms. Though the number of unique tRFs (143) expressed in the sperm was 16 times lower than piRNA (2290) and 2.8 times lower than miRNA (401), their expression levels were relatively high, making them the most abundant sncRNA expressed in male gametes. tRF5b and tRF5c were the dominant tRNA derivatives, that have been previously reported to be involved in epigenetic inheritance (Kumar et al. 2016). We also detected tRF3a, b and c forms that not only have lengths comparable to mature miRNAs, but are also known to function like miRNAs and follow similar target binding rules. The most highly expressed tRFs included Gly-GCC, Glu-CTC, Glu-TTC, Val-CAC and His-GTG, which were also reported to be the most abundant tRFs in mouse model systems (Chen et al. 2016; Sharma et al. 2016). tRF-Gly-GCC has been shown to repress expression of genes regulated by endogenous retroelement MERVL in early embryo development. Our study also identified low Gly-GCC expression to be associated with low sperm concentration, though we also found 5 new tRFs to contribute to this association. Among the differentially expressed tRFs, Cys-GCA was previously reported to be downregulated in a high-fat diet mouse model (Chen et al. 2016).

The effectiveness of the 6 weeks dietary intervention was evident by a significant increase in the vitamin D levels of serum, and the EPA and DHA percentage in RBCs of the participants in the intervention group. 7/15 differentially expressed miRNAs were found to target genes enriched in fatty acid biosynthesis and metabolism, demonstrating the potential of the dietary intervention to generate a physiological change sufficient to be signaled to the male germline and be epigenetically captured by alterations to sncRNA. Around 18.4% (422/2290) of baseline piRNAs and 10.7% (12/112) of diet induced piRNAs were expressed from transposable elements (TE) such as LINES, SINES and LTRs. Since one of the functions of piRNAs is to protect genome integrity during spermatogenesis by silencing transposable elements, a number of them were found to originate directly from TE transcripts in the sense orientation. piRNAs are also known to have non TE related functions, such as regulation of mRNA and lincRNA expression through post-transcriptional gene silencing (Larriba and Del Mazo 2018). We found genes related to baseline piRNAs to be enriched in pathways such as Golgi and Endomembrane system organization. Strikingly, we also found 17 baseline piRNAs to originate from tRNAs, suggesting the existence of a regulatory cross-talk between the two. Further in-depth studies are warranted to develop deeper insights into sperm sncRNA regulation.

This study has some limitations. To avoid small sample size errors and detection of false positives, we restricted the baseline analysis to the high confidence sncRNAs detected in all 17 participants. This high stringency analysis may have limited the sensitivity of detection and missed some sncRNAs. Also, due to the limited functional characterization of the downstream targets of human tRFs, we were unable to reflect on the regulatory outcomes of their altered expression upon diet intervention. However, the high-resolution tRF and piRNA mapping attempted in this study, unfolded the not yet known complexity in the expression of their subtypes in human sperms. Expression of a subset of piRNA from the tRNA locus also identified the potential cross-talk between different sncRNA, that might be essential to effectively capture the effects of external exposures in the male germline.

In conclusion, the epigenetic programing of male gametes is highly labile and sensitive to alterations in lifestyle, such as diet. Effects of short-term dietary exposures can be captured as “environmental messages” in the form of altered sncRNA expression profiles in the male germline, and be likely transmitted to the next generation if these changes occur around conception. These studies suggest that paternal diet may have long lasting effects on the health of the offspring and consumption of a healthy diet in the preconception period may help enhance the quality of sperm by epigenetic coding of the sncRNAs conferring beneficial health effects to the next generation.

## Methods

### Sample information and study design

The PREPARE trial is a single center, randomized controlled trial, in which 111 men and their partners about to undergo a cycle of IVF treatment provided written informed consent to be recruited to the study (Kermack et al. 2014). Ethical approval for this study was granted by the regional Research Ethics Committee (13/SC/0544). Exclusion criteria included eating oily fish (as defined by the UK Food Standards Agency) more than once a week, previously diagnosed diabetes, and any medical contraindication to the dietary supplements. Demographic data from the men and information on their diet and physical activity were collected on a standard study proforma at the time of recruitment and their BMI was determined. The mean age of the participants was 36 (± 5.5) years and mean BMI was 27.0 (± 4.0) kg/m^2^ (**Supplemental File S1**). Study participants were randomized into the control and intervention diet groups. The intervention group were given a daily fruit-based dietary supplement drink containing 2 grams DHA and EPA (1.2 g DHA and 0.8 g EPA) and 10 micrograms vitamin D (Nutrifriend 2000, Smartfish, Oslo, Norway) and olive oil and olive oil spread to use as part of their diet, while the control group received a placebo drink (the same fruit-based drink but without omega-3 fatty acids or vitamin D (Smartfish, Oslo, Norway)) and sunflower oil and sunflower oil based spread to use as part of their diet. The olive oil, sunflower oil and both spreads were purchased from a local supermarket and were repackaged for use in the study. The drinks, oils and spreads were all provided in unmarked containers and the trial was double blind. The duration of intervention was 6 weeks. 102 participants completed the trial and had semen samples collected at both pre and post-intervention visits. sncRNAseq profiling was performed on a subset of these participants (N=17; 9 from the control group and 8 from the intervention group) (**Figure 1A**).

### Semen analysis

Semen analysis was performed by a blinded andrologist on all samples in accordance with the WHO guidelines (Cooper and Castilla 2009). Sample liquefaction and viscosity; the volume of semen produced; sperm concentration; percentage of motile and progressively motile sperm; and percentage of sperm with normal morphology were recorded. The sample was considered normal if at least 1.5 ml of semen was produced; it liquefied within one hour and was not viscous; and it had a least 15 million spermatozoa per ml of ejaculate with a progressive motility of at least 32% and a normal morphology of at least 4%.

Semen (2 ml or the maximum volume available if less) was placed on a density gradient (created using 1 ml of Nidacon PureSperm®100 90% solution overlaid with a 1ml of Nicadon PureSperm®100 45% solution) and centrifuged at 400 g for 20 minutes to separate the seminal plasma from the sperm. The supernatant (seminal plasma) was removed and stored in 1 ml aliquots at -80°C. The sperm pellet was washed with phosphate-buffered saline (PBS) and centrifuged at 400 g for a further 5 minutes, this sperm pellet was collected and stored at -80°C prior to analysis. For DNA fragmentation analysis, semen was left to liquefy for at least 30 minutes following ejaculation. Between 100 µl and 200 µl was then placed into a tube and plunged into liquid nitrogen to snap freeze and then stored at -80°C until analysis. Sperm DNA fragmentation was assessed using the SpermComet assay, a second generation sperm DNA test (Lewis et al. 2013).

### Blood sampling

Blood was collected into lithium heparin and serum separating tubes at the time of recruitment and on the day of partner’s oocyte retrieval. Blood collected into lithium heparin was centrifuged at 1000 g for 15 minutes at 20°C. The plasma was removed and the red cell pellet was washed in PBS and centrifuged at 400 g for 10 minutes at room temperature. The wash step was repeated twice. The red cell pellet was then stored as a 0.5 ml aliquot at -80°C prior to fatty acid analysis. Blood collected into SST was mixed and centrifuged at 1500 g for 10 minutes. The supernatant (serum) was removed and stored in aliquots at -80°C prior to vitamin D analysis.

### Fatty acid analysis

Fatty acid analysis was performed on red blood cells, seminal plasma and sperm. In all cases, total lipid was extracted into chloroform: methanol (2:1, vol/vol). The lipid extract was heated to 50°C for 2 hours with 2% methanol in sulphuric acid to produce fatty acid methyl esters (FAMEs). FAMEs were separated and identified by gas chromatography performed according to conditions described elsewhere (Fisk et al. 2014). FAMEs were identified by comparison of run times with those of authentic standards. Fatty acid concentrations are expressed as % of the total fatty acids present. Analysis on the sperm was only performed if the sperm concentration was greater than 15 million per ml and there was enough semen remaining following clinical use on the day of oocyte retrieval.

### Vitamin D analysis

Serum vitamin D concentrations were determined by Liquid Chromatography/Tandem Mass Spectrometry (Waters, Milford, MA, USA) in the Chemical Pathology Laboratory, University Hospital Southampton NHS Foundation Trust, a member of the Vitamin D EQA scheme (DEQAS).

### Semen sample processing and RNA extraction

Total RNA including sncRNA was extracted from frozen sperm pellets using the RNeasy Micro Kit (Qiagen, UK) with a modified protocol adapted from Goodrich, *et al*., (Goodrich et al. 2013) which was optimised for recovery of sperm RNA. Briefly, sperm pellets were thawed, mixed with Qiazol (Qiagen, UK) and manually disrupted. Chloroform was added, samples incubated (room temperature for 5 minutes) and centrifuged (12000 *x g*) for 20 minutes. The upper aqueous layer was removed and mixed with ethanol. Samples were added to micro RNeasy spin columns (Qiagen, UK) and centrifuged for 15 seconds (8000 *x g*). Spin columns were washed using supplied buffers and ethanol. Samples were transferred to RNAse free collection tubes and total RNA eluted. RNA concentrations were quantified using Qubit total and microRNA assays (Thermo Fisher, UK), RNA was snap frozen on dry ice and stored at -80 degrees.

### sncRNA sequencing

Sperm RNA samples were selected from both control and dietary intervention groups based on the quantity of RNA post extraction (n=17). 10µl of RNA was used to perform small RNA-seq (∼15 million reads) using the smallRNA workflow (Oxford Genomics, OGC). NEBNext Small RNA Library Prep Set for Illumina (E7330) was used as per manufacturers’ instructions. Briefly, 3’ SR adapter ligation was carried out followed by RT primer hybridisation to the mRNA. Denatured 5’ SR adapter was then ligated, and first strand cDNA reverse transcription was performed. After second strand synthesis with indexed (TruSeq-based single indexed i7) and P5 primers, dsDNA was then cleaned-up using Ampure XP beads, and the libraries were size selected by Pippin (125-160bp). The concentration of the small RNA libraries was determined by Picogreen before multiplexing for sequencing. The final size distribution of the pooled library was determined using Agilent Tapestation, and quantified by Qubit (Thermofisher, UK), before sequencing on the HiSeq2500 (Illumina, UK) (2X runs) as 50bp single reads. Data were aligned to the reference genome and FASTQ files generated.

### Processing of sncRNA sequencing data

FastQC (https://www.bioinformatics.babraham.ac.uk/projects/fastqc/) was used to perform quality check (QC) on the sequencing reads. All samples passed the mean per base sequence quality scores ≥30. Average length of sequenced reads was 51nt, inclusive of the adaptors. Trimmomatic 0.38 (Bolger et al. 2014) was used to remove the adapter sequences, and sequence reads < 10nt were excluded from the subsequent analysis. exceRpt small RNA-seq analysis pipeline available under the Genboree workbench (https://www.genboree.org/site/) was used to profile and assess the relative abundance of sncRNA subtypes. Since the current version of Genboree does not include the latest release for miRNA and tRF databases, we additionally used specific annotator tools to report the final results. Mapper.pl and quantifier.pl modules of the miRDeep2 tool (Friedlander et al. 2012) and the latest miRBase (http://mirbase.org/) release 22 were used to annotate and quantify the known human miRNA.

Following parameters were used for the mapper and quantifier module taking sample 005mv1-249 as an example:

*mapper*.*pl 005mv1-249-trim*.*fastq -e -g 249 -h -i -j -l 15 -m -s 005mv1-249-trim-col*.*fa –v quantifier*.*pl -p hsa-precursor-miRNAs-R22*.*fa -m hsa-mature-miRNAs-R22*.*fa -r 005mv1-249-trim-col*.*fa -t hsa*

For tRF analysis, human tRNA sequences were downloaded from the tRNA database (http://gtrnadb.ucsc.edu/). For each tRNA gene, the DNA sequences extracted from the genome included additional 100 bases upstream and 200 bases downstream of the mature tRNA. Since ‘CCA’ is added to the 3’ end of the tRNA by tRNA nucleotidyltransferase during the tRNA maturation process, this triplet nucleotide sequence was also included at the 3’ end of the extracted DNA sequences to avoid sequence mismatch during the mapping process. The sperm sncRNA sequencing reads were then mapped to the extracted tRNA sequence by using the BLASTn tool. Sequence alignments with ≤1 nt mismatch were retained for subsequent analysis. The blast output file was parsed to get information on the mapped position of small RNA on the tRNA genes and segregate the sequences into specific subtypes: 5’ tiRs, 5’ (a,b,c), 3’ (a,b,c) and 1-tRFS.

For piRNA expression analysis, Genboree platform was used to quantify and profile the human piRNAs from the piRNABank.

### sncRNA association with paternal factors

For baseline characterization of sncRNA, raw QCed reads were processed to generate expression values in counts per million (cpm). sncRNA with cpm of ≥1 in all 17 subjects in the pre-intervention group were retained and normalized using Trimmed Mean of M (TMM) method. The normalized data was further subjected to voom with sample specific quality weights transformation to down weigh the outlier samples. Limma Bioconductor package (Ritchie et al. 2015) was used for association analysis of baseline sncRNA with the paternal factors such as age, BMI, alcohol/caffeine consumption and sperm quality. The moderated t-statistics of association was computed by empirical Bayes (eBayes), using a linear model fit (Lmfit). Associations with FDR corrected p value < 0.05 were reported as significant changes in response to a unit change in the assessed paternal factor.

### Differential expression of sncRNA between the control and intervention groups

Differentially expressed sncRNA between the control and intervention groups were analyzed using the Limma Bioconductor package (Ritchie et al. 2015). sncRNAs that had a cpm of ≥1 in at least 8 samples within each group were retained and normalized using the Trimmed Mean of M (TMM) method. The normalized data was subjected to double voom with sample specific quality weights transformation to down weigh the outlier samples. Since the samples were collected at two time points from the same individual, a paired analysis was performed using Limma’s duplicate correlation method and blocking the ID. This analysis was adjusted for the confounding effects of age, BMI, sperm concentration and motility. The moderated t-statistics of differential expression was computed by empirical Bayes (eBayes), using a linear model fit (Lmfit) and comparing the overall effect of intervention over control using the following equation: (Intervention.T2 - Intervention.T1) - (Control.T2 - Control.T1), where T2 is the time point after the 6 weeks of intervention and T1 is the time point of the first visit. Since none of the sncRNAs passed the FDR corrected p-value of ≤ 0.05, we reported the sncRNA that passed the unadjusted p-value cut-off of ≤ 0.01 and log2(1.5) fold change.

### Determining the biological significance of sncRNAs

#### miRNA target information and pathway enrichment using DIANA miRPath and IPA

DIANA-mirPath v3.0 (http://snf-515788.vm.okeanos.grnet.gr/) is an online software suite that makes use of the predicted miRNA targets provided by the DIANA-microT-CDS algorithm or experimentally validated miRNA-target interactions obtained from DIANA-TarBase. These predicted and/or experimentally validated targets are subsequently used for KEGG and Gene Ontology analysis. This software suite was used for the evaluation of miRNA target genes and the related functional pathways.

In addition to DIANA-mirPath we also used IPA (QIAGEN Inc., https://www.qiagenbioinformatics.com/products/ingenuity-pathway-analysis) to assess the functional roles of miRNAs. This tool derives miRNA target information from databases such as TarBase, miRecords and TargetScan. Experimentally known as well as high confidence (high scoring) targets were filtered and genes targeted by ≥2 miRNAs were used for the core analysis in IPA that reports the significantly enriched pathways. A threshold –log10 p-value of 2 was used to report the pathways in which the target genes were significantly enriched.

#### Determining the targets of the baseline and differentially expressed piRNAs

Two approaches were used to determine the functional annotation of piRNAs identified in this study. The first approach involved annotation of the genomic coordinates of the DE-piRNAs and baseline piRNAs using the annotatePeaks.pl program in the Homer motif analysis software package (http://homer.ucsd.edu/homer/). The second approach involved comparison of the piRNAs (piRNABank) (Sai Lakshmi and Agrawal 2008) with the piRBase database (Wang et al. 2019) that comprises of information on repeat related and genes related piRNAs.

### Data access

All raw and processed sncRNA sequencing data generated in this study have been submitted to the NCBI Gene Expression Omnibus (GEO; https://www.ncbi.nlm.nih.gov/geo/) under the accession number GSE159752. Due to ethical concerns, supporting clinical data cannot be made openly available. However, the PREPARE study team can provide the data upon request subject to appropriate approvals after a formal application to the PREPARE Study Oversight Group through the corresponding author.

## Supporting information

Combined-Supplemental-Figures-Tables

S1-Demographics

S2-Baseline-sncRNA

S3-Baseline-tRF-seq

S4-Baseline-piRNA-Homer-annot

S5-Baseline-piRNA-piRBase-annot

S6-pre-int-factors-miRNA

S7-DE-sncRNA

S8-DE-piRNA-Homer-annot

S9-DE-piRNA-piRBase-annot

## Competing interests

NK, KMG, KAL and YSC are part of an academic consortium that has received research funding from Abbott Nutrition, Nestec, BenevolentAI Bio Ltd. and Danone. PCC serves on the Scientific Advisory Board of Smartfish, the producer of the drinks used in the study. PCC has received advisory and/or speaking honoraria from Fresenius-Kabi, B. Braun, Baxter Healthcare, Abbott Nutrition, and Danone/Nutricia, sellers of parenteral and enteral feeds, and from Pronova BioPharma/BASF AS and Smartfish, sellers of products containing omega-3 fatty acids. KMG has received reimbursement for speaking at conferences sponsored by companies selling nutritional products. The remaining authors declare no competing interests.

Trial registration number: ISRCTN50956936

Trial registration date: 10/02/2014

## Acknowledgements

We wish to thank the study participants from Complete Fertility, Southampton for making this project possible as well as the staff at the institutions for assistance in project management. Sperm sncRNA analysis was funded by EpiGen Global Research Consortium Seed Funds available to NK and KAL. Additional funds for sncRNA data analysis in Singapore was supported by the Strategic Positioning Fund and IAFpp funds (H17/01/a0/005) available to NK through Agency for Science, Technology and Research (A*STAR), Singapore (award number SPF 002/2013). We also thank the Oxford Genomics Centre at the Wellcome Centre for Human Genetics (funded by Wellcome Trust grant reference 203141/Z/16/Z) for the generation of the sequencing data. The PREPARE trial was funded by the NIHR Southampton Biomedical Research Centre. The drinks (both intervention and control) were designed, manufactured and supplied by Smartfish. KMG is supported by the UK Medical Research Council (MC_UU_20/4), the US National Institute On Aging of the National Institutes of Health (award number U24AG047867), the UK Economic and Social Research Council and the Biotechnology and Biological Sciences Research Council (award number ES/M0099X/), the National Institute for Health Research (as an NIHR Senior Investigator (NF-SI-055–0042) and through the NIHR Southampton Biomedical Research Centre), the British Heart Foundation (RG/15/17/3174) and the European Union’s Erasmus□+□Capacity-Building Program (ImpENSA, grant agreement number 598488-EPP-1-2018-1-DE-EPPKA2-CBHE-JP). KAL is supported by the British Heart Foundation (RG/15/17/3174), Diabetes UK (16/0005454). The funding bodies had no role in the design, collection, analysis, and interpretation of data, writing of the paper, or decision to submit for publication.

## Author contributions

NK, KAL and KMG conceptualized the study and NK and KAL supervised the sncRNA study in Singapore and UK, respectively. AJK, PCC, FDH and NSM designed their PREPARE trial, AJK, KD and SJW contributed to trial sample and data collection and TPXY, HLF and SL to sample analysis. MB processed the sperm samples for RNA extraction and sncRNA sequencing, CV, PFT, PK, AD and NK processed, analyzed, and interpreted the sncRNA sequencing data. CV, PK, KAL, KMG, MB and NK contributed to writing the initial draft of the manuscript. JH, CV, PFT and NK developed the iDad_db database. All authors critically reviewed the manuscript for intellectual and scientific content and approved the final manuscript.

## Supplemental Figures and Table Legends

**Supplemental Figure S1. Chromosome-wise enrichment analysis of baseline miRNAs in sperms**. Horizontal dashed lines indicate the z-score cut-offs for over (≥ 2) and under (≤ -2) representation of miRNAs.

**Supplemental Figure S2. Chromosome-wise enrichment analysis of baseline tRFs in sperms**. Horizontal dashed lines indicate the z-score cut-offs for over (≥ 2) and under (≤ -2) representation of tRFs.

**Supplemental Figure S3. Chromosome-wise enrichment analysis of baseline piRNAs in sperms**. Horizontal dashed lines indicate the z-score cut-offs for over (≥ 2) and under (≤ -2) representation of piRNA

**Supplemental Figure S4. IPA based pathway enrichment analysis of miRNA target genes associated with age**.

Highlighted in purple are the known ageing related gene pathways. The dotted red line represents the ratio of the miRNA target genes to the total number of genes known to be present in that pathway.

**Supplemental Figure S5. IPA based pathway enrichment analysis of miRNA target genes associated with sperm concentration**.

Highlighted in purple are the known sperm concentration related gene pathways. The dotted red line represents the ratio of the miRNA target genes to the total number of genes known to be present in that pathway.

**Supplemental Table S1:** Top 10 pathways (FDR q-value < 0.05) enriched in 234 genes associated with baseline piRNA listed in Supplemental File S5, sheet 6.

**Supplemental Table S2**. Baseline tRFs and piRNAs associated with BMI and sperm concentration at -log10 adjusted p-value ≥ 1.3

**Supplemental Table S3. Diet intervention induced alterations in sncRNA expression**

(A) The differentially expressed miRNAs (intervention vs. control) identified by comparing miRDeep2 derived miRNA expression profile. This analysis was adjusted for age, BMI, sperm concentration and sperm motility. miRNAs with ≥ log2(1.5) fold change and an unadjusted -log10 p-value ≥2 are reported.

(B) Differentially expressed tRFs identified after adjusting for age, BMI, sperm concentration and sperm motility using log2(1.5) fold change cut-off and an un-adjusted -log10 p-value ≥2.

**Supplemental Table S4. Pathway analysis of miRNA targeted genes altered by diet intervention**.

(A) Gene pathways identified using DIANA mirPath software, at a FDR corrected p-value of <0.05.

(B) mirPath micro-T CDS target prediction scores for 7 DE-miRNAs targeting fatty acid metabolism genes.

## Supplemental Files

**Supplemental File S1. Subject characteristics and demographics, sperm analysis, semen analysis and blood measures of the study participants**

Sheet 1: sncRNA sub set (N=17) pre-intervention

Sheet 2: Full set (N=102) pre-intervention

Sheet 3: sncRNA sub set (N=17) post-intervention

Sheet 4: Full set (N=102) post-intervention

**Supplemental File S2. Pre-intervention baseline characterization of sncRNA landscape**

Sheet 1: Expression profile of 401 mature miRNAs

Sheet 2: Expression profile of 143 tRFs

Sheet 3: Expression profile of 2290 piRNAs

**Supplemental File S3. Baseline tRF sequences present in the pre-intervention samples**

Sheet 1: Most abundant sequence for each tRF ID across the 17 samples.

Sheet 2: Representative tRF sequence for each tRF ID

**Supplemental File S4. Homer annotation of all the genomic locations of baseline piRNAs**

Sheet 1: Baseline piRNAs overlapping with transposable elements (TEs)

Sheet 2: Baseline piRNAs overlapping with tRNAs

Sheet 3: Baseline piRNAs not overlapping with transposable elements (NTEs)

**Supplemental File S5. Mapping of piRNABank identified baseline piRNA to piRBase annotations**

Sheet 1: piRNABank piRNA identifiers that mapped to piRBase

Sheet 2: LINE related baseline piRNAs in piRBase

Sheet 3: SINE related baseline piRNAs in piRBase

Sheet 4: LTR related baseline piRNAs in piRBase

Sheet 5: Satellite related baseline piRNAs in piRBase

Sheet 6: Genes related baseline piRNAs in piRBase

**Supplemental File S6. miRNA associated with age and sperm concentration**

Sheet 1: miRNAs associated with age (adjusted –log10 p value ≥ 1.3)

Sheet 2: miRNAs associated with sperm concentration (adjusted –log10 p value ≥ 1.3)

**Supplemental File S7. Differentially expressed sncRNA between control and intervention** sncRNA highlighted in purple represent the DE-sncRNA identified after adjusting for the effects of age, BMI, sperm concentration and motility.

Sheet 1: Differentially expressed miRNA (unadjusted –log10 p value ≥ 2).

Sheet 2: Differentially expressed tRF (unadjusted –log10 p value ≥ 2).

Sheet 3: Differentially expressed piRNA (unadjusted –log10 p value ≥ 2).

**Supplemental File S8: Homer annotation of all the genomic locations of DE-piRNAs**

Sheet 1: DE-piRNAs overlapping with transposable elements (TEs)

Sheet 2: DE-piRNAs not overlapping with transposable elements (NTEs)

**Supplemental File S9: Mapping of piRNABank identified DE-piRNA to piRBase annotations**

Sheet 1: piRNABank piRNA identifiers that mapped with piRBase.

Sheet 2: Repeats related DE-piRNAs in piRBase

Sheet 3: Genes related DE-piRNAs in piRBase

