## Supplementary material for "Short-term diet intervention alters the small non-coding RNA (sncRNA) landscape of human sperm": Combined-Supplemental-Figures-Tables

### and * represent joint authorships

[1]Singapore Institute for Clinical Sciences (SICS), A*STAR, Brenner Centre for Molecular Medicine, Singapore.
[2]School of Human Development and Health, Faculty of Medicine, University of Southampton, Southampton, UK.
[3]NIHR Southampton Biomedical Research Centre, University Hospital Southampton, NHS Foundation Trust and University of Southampton, Southampton, UK.
[4]Complete Fertility, Princess Anne Hospital, Southampton, UK
[5]University Hospital Southampton NHS Foundation Trust, Southampton, UK.
[6]Queen's University, Belfast, Northern Ireland, UK.
[7]Examen Lab Ltd, Belfast, Northern Ireland, UK
[8]Department of Obstetrics and Gynaecology, Yong Loo Lin School of Medicine, National University of Singapore, Singapore.
[9]Liggins Institute, University of Auckland, Auckland, New Zealand.
[10]London Women's Clinic, London UK.
[11]Department of Biochemistry and Molecular Genetics, University of Virginia, School of Medicine, Charlottesville, VA, USA.
[12]MRC Lifecourse Epidemiology Unit, University of Southampton, Southampton, UK.
[13]School of Biological Sciences, University of Southampton, Southampton, UK.
[14]Department of Biochemistry, Yong Loo Lin School of Medicine, National University of Singapore (NUS), Singapore.

**This PDF file includes:**

- Supplemental Figures S1 - S5
- Supplemental Tables S1 - S4
- List of Supplemental Files S1 - S9

**Supplemental Figure S1. Chromosome-wise enrichment analysis of baseline miRNAs in sperms**

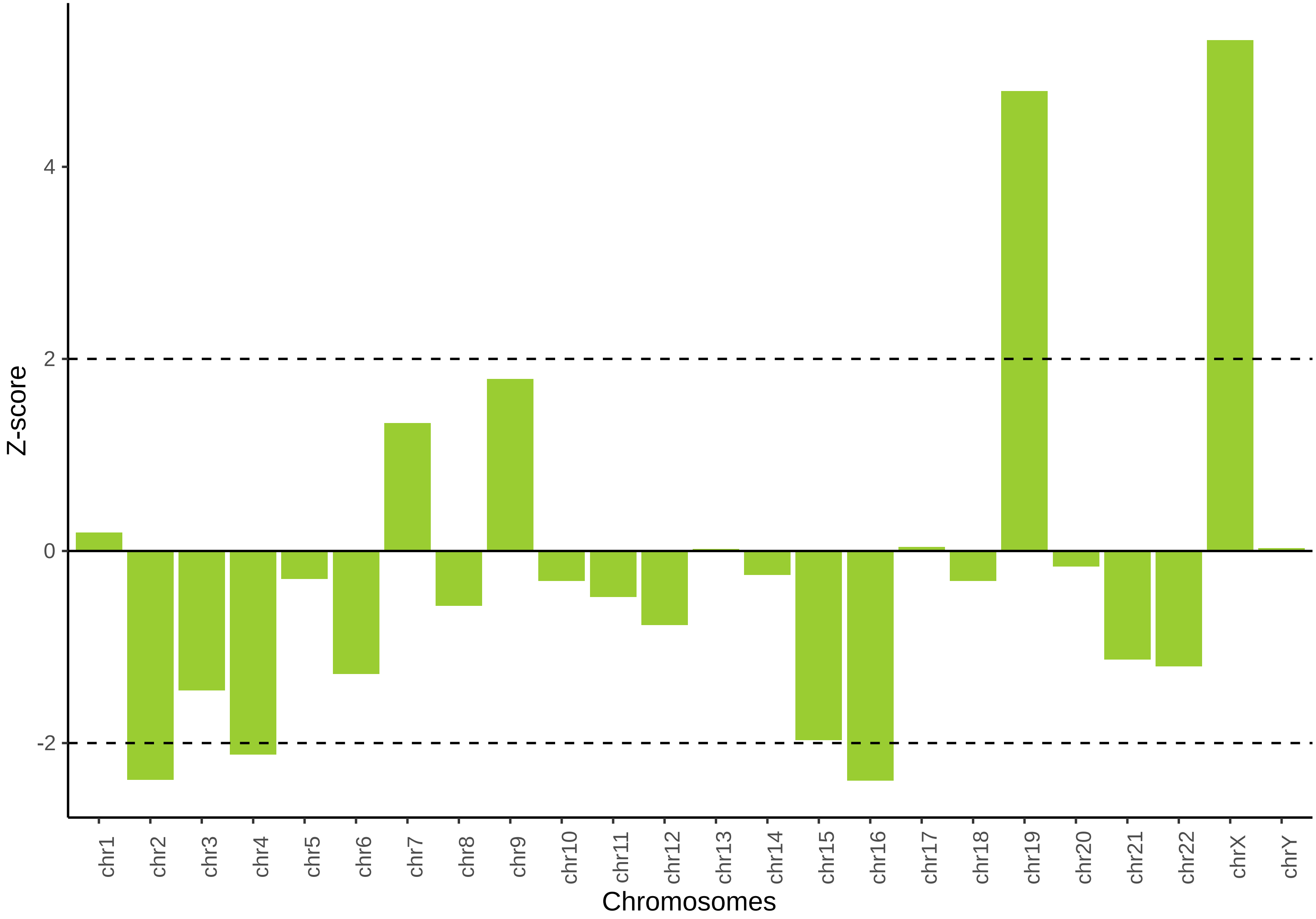


Horizontal dashed lines indicate the z-score cut-offs for over (≥ 2) and under (≤ -2) representation of miRNAs.

**Supplemental Figure S2. Chromosome-wise enrichment analysis of baseline tRFs in sperms**

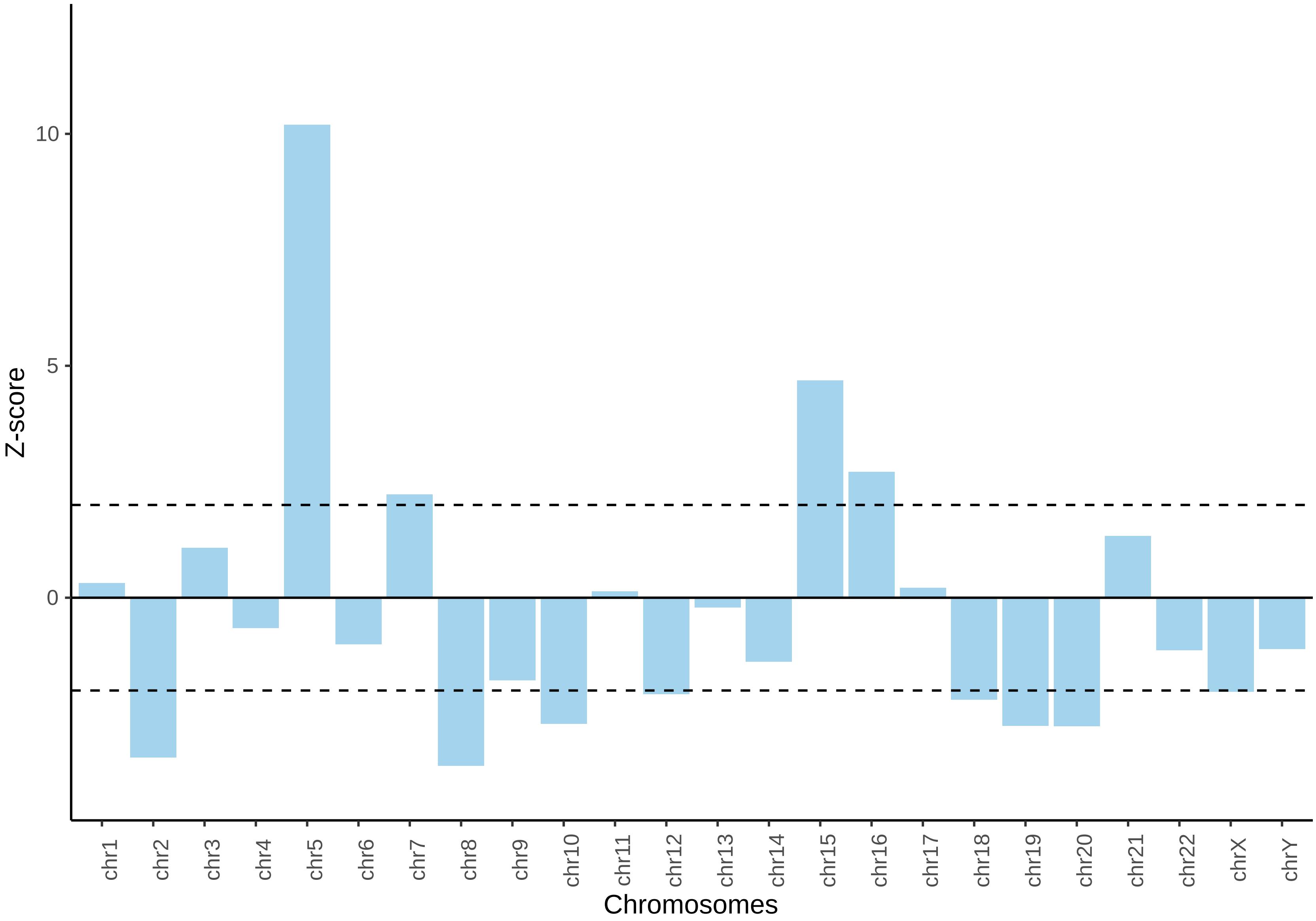


Horizontal dashed lines indicate the z-score cut-offs for over (≥ 2) and under (≤ -2) representation of tRFs.

**Supplemental Figure S3. Chromosome-wise enrichment analysis of baseline piRNAs in sperms**

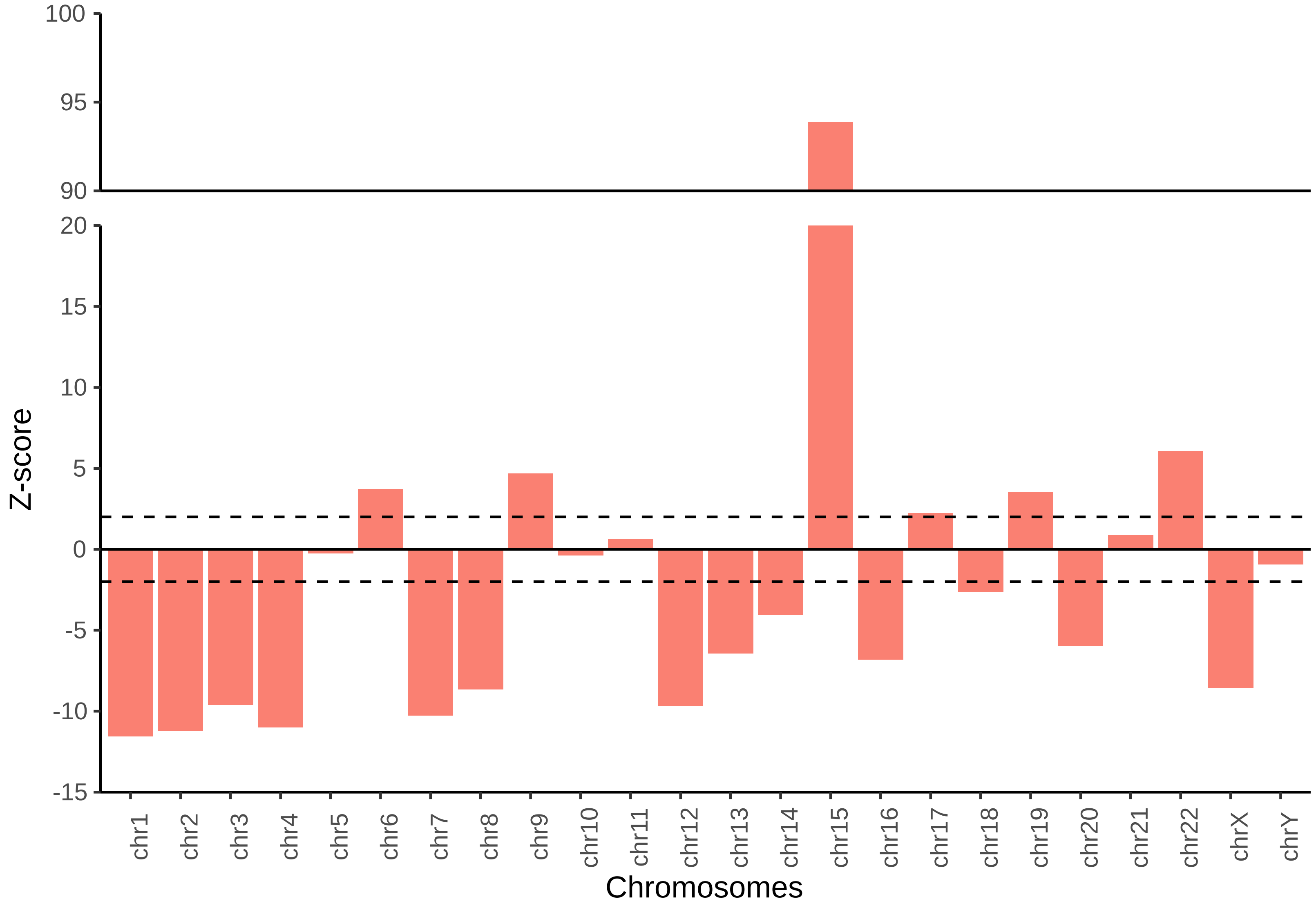


Horizontal dashed lines indicate the z-score cut-offs for over (≥ 2) and under (≤ -2) representation of piRNA.

**Supplemental Figure S4. IPA based pathway enrichment analysis of miRNA target genes associated with age**

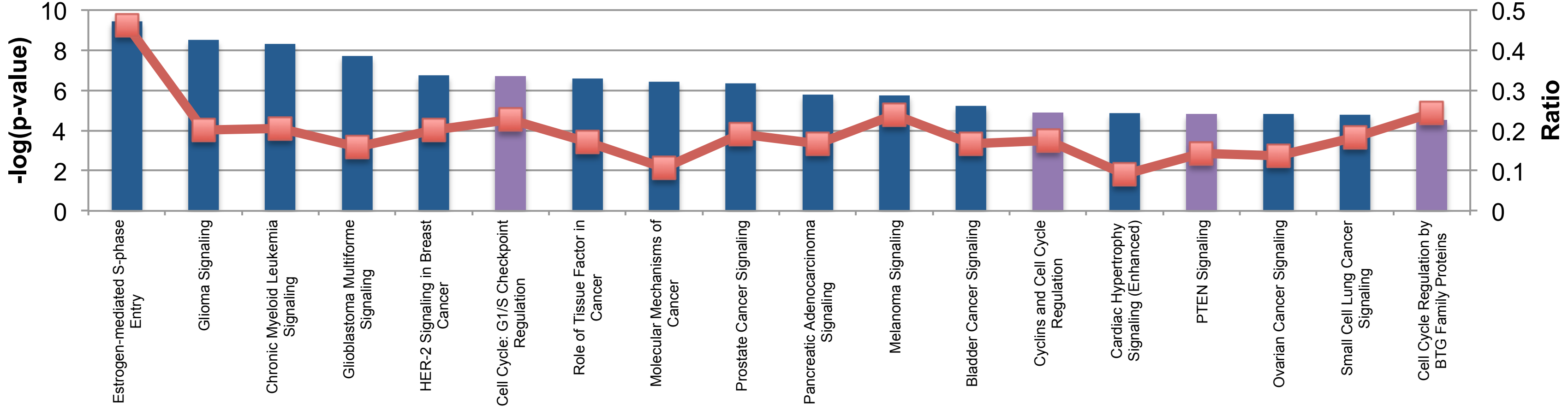


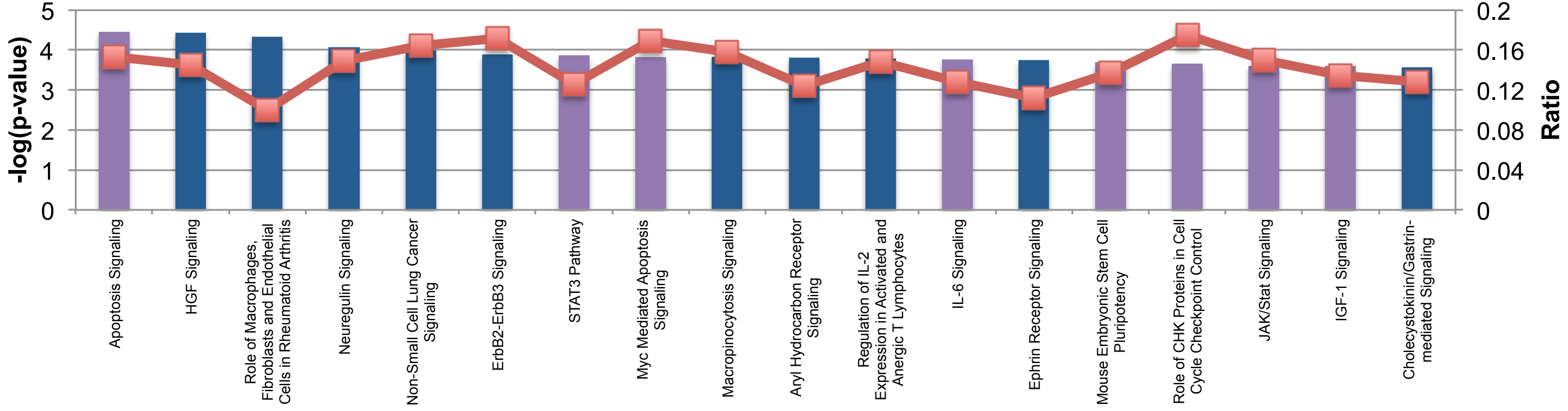

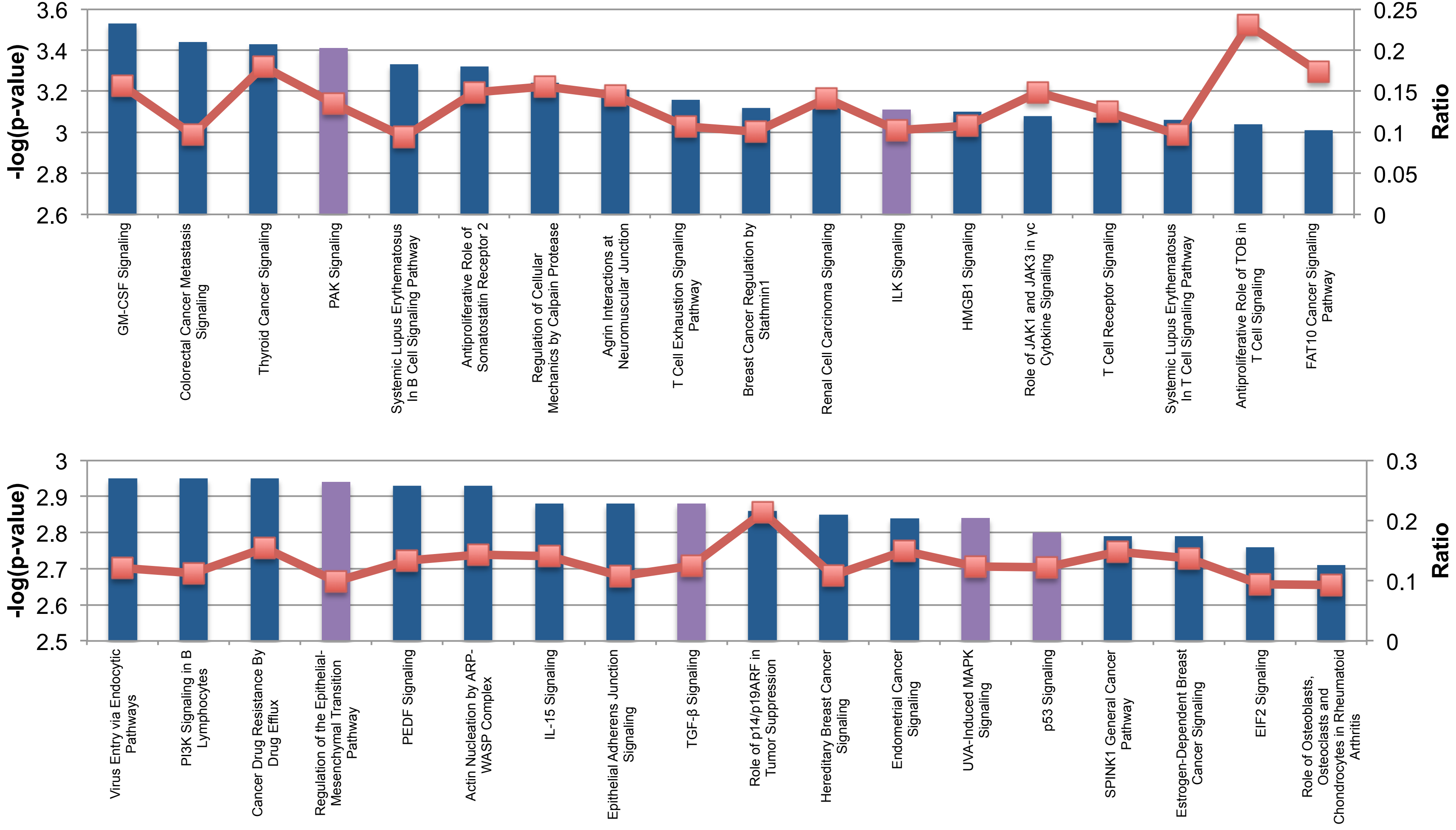

-log(p-value)
Ratio
3.6
3.4
3.2
3
2.8
2.6
0.25
0.2
0.15
0.1
0.05
0
GM-CSF Signaling
Colorectal Cancer Metastasis Signaling
Thyroid Cancer Signaling
PAK Signaling
Systemic Lupus Erythematosus In B Cell Signaling Pathway
Antiproliferative Role of Somatostatin Receptor 2
Regulation of Cellular Mechanics by Calpain Protease
Agrin Interactions at Neuromuscular Junction
T Cell Exhaustion Signaling Pathway
Breast Cancer Regulation by Stathmin1
Renal Cell Carcinoma Signaling
ILK Signaling
HMGB1 Signaling
Role of JAK1 and JAK3 in γc Cytokine Signaling
T Cell Receptor Signaling
Systemic Lupus Erythematosus In T Cell Signaling Pathway
Antiproliferative Role of TOB in T Cell Signaling
FAT10 Cancer Signaling Pathway
-log(p-value)
Ratio
3
2.9
2.8
2.7
2.6
2.5
0.3
0.2
0.1
0
Virus Entry via Endocytic Pathways
PI3K Signaling in B Lymphocytes
Cancer Drug Resistance By Drug Efflux
Regulation of the Epithelial-Mesenchymal Transition Pathway
PEDF Signaling
Actin Nucleation by ARP-WASP Complex
IL-15 Signaling
Epithelial Adherens Junction Signaling
TGF-β Signaling
Role of p14/p19ARF in Tumor Suppression
Hereditary Breast Cancer Signaling
Endometrial Cancer Signaling
UVA-Induced MAPK Signaling
p53 Signaling
SPINK1 General Cancer Pathway
Estrogen-Dependent Breast Cancer Signaling
EIF2 Signaling
Role of Osteoblasts, Osteoclasts and Chondrocytes in Rheumatoid Arthritis

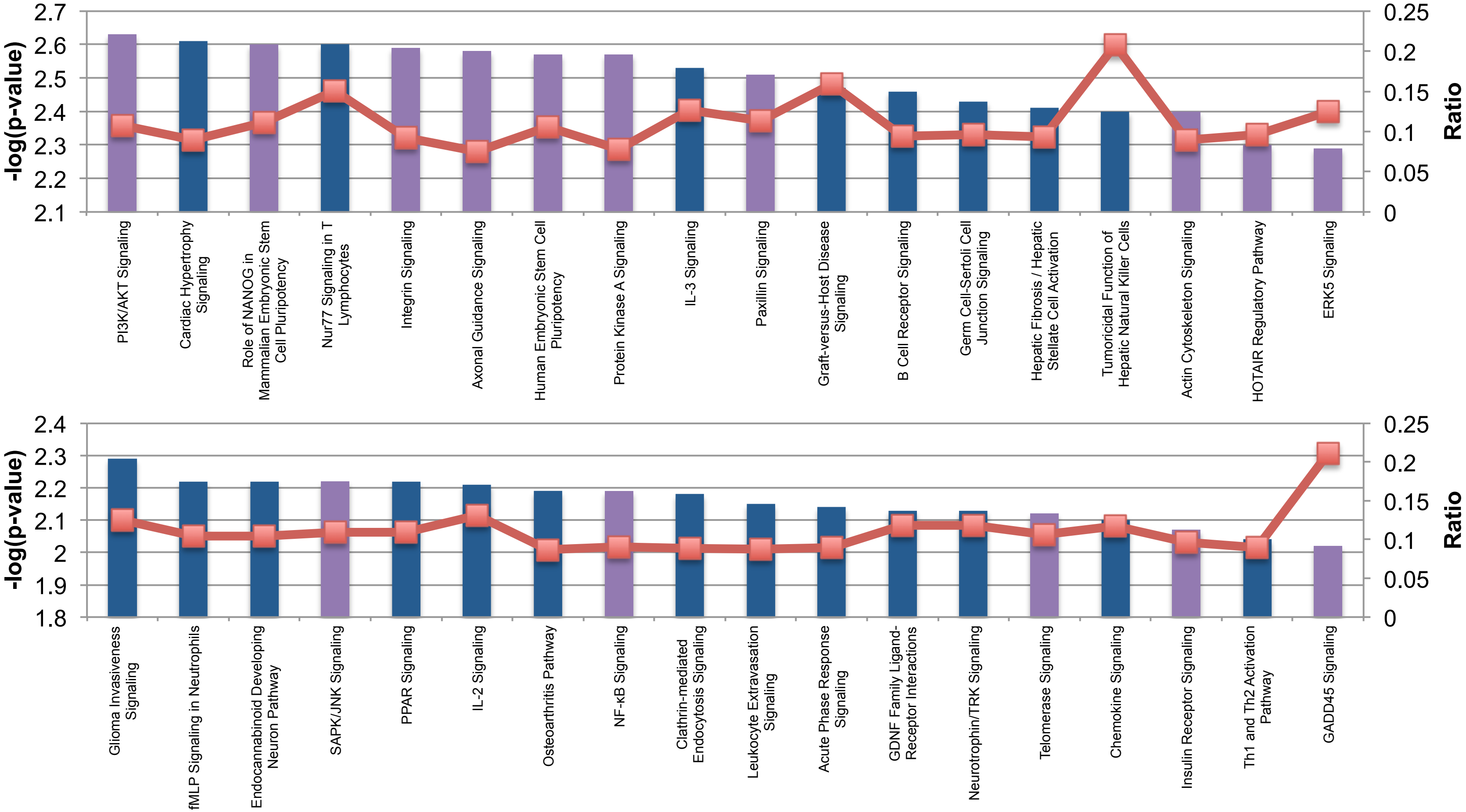


Highlighted in purple are the known ageing related gene pathways.
The dotted red line represents the ratio of the miRNA target genes to the total number of genes known to be present in that pathway.

**Supplemental Figure S5. IPA based pathway enrichment analysis of miRNA target genes associated with sperm concentration**

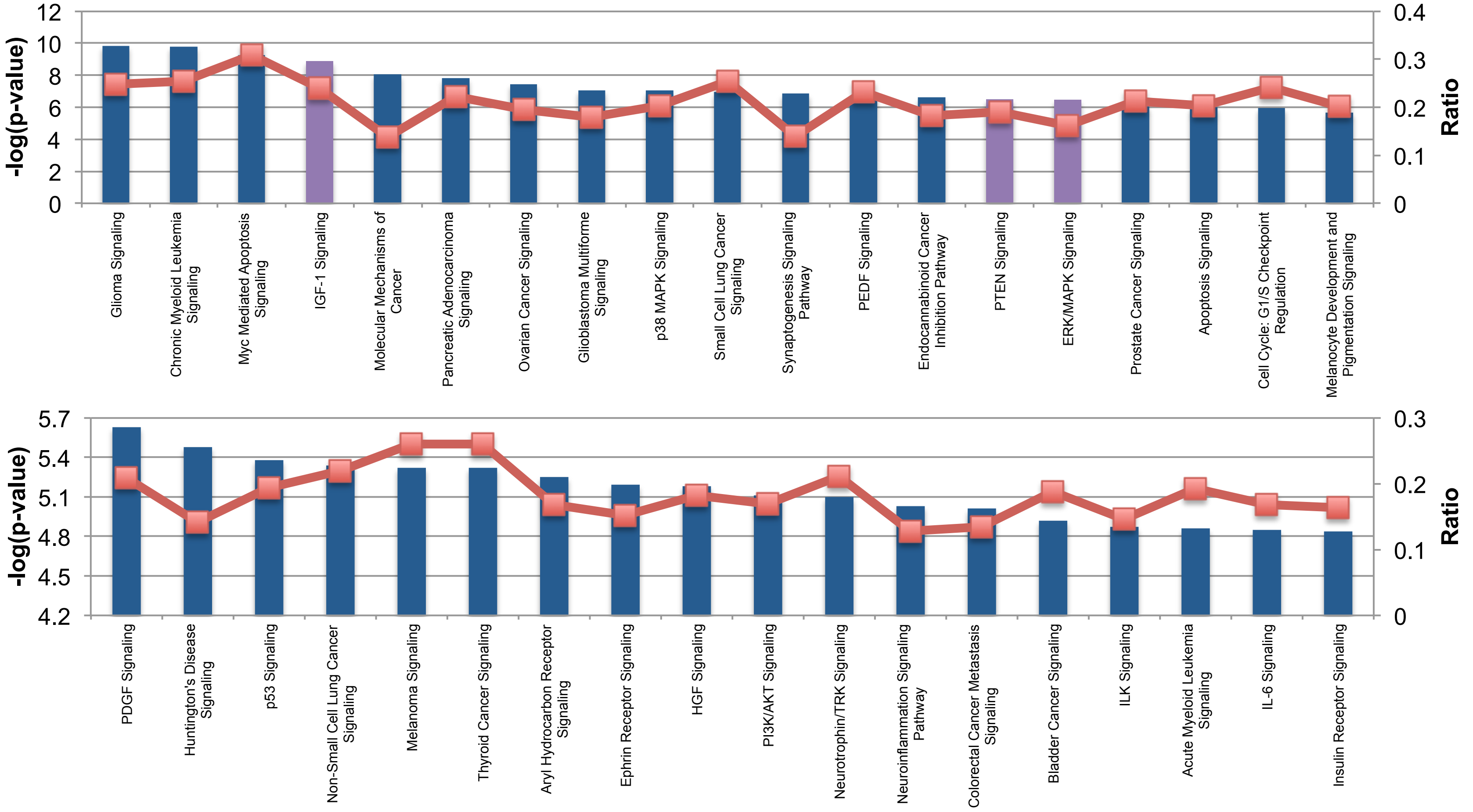

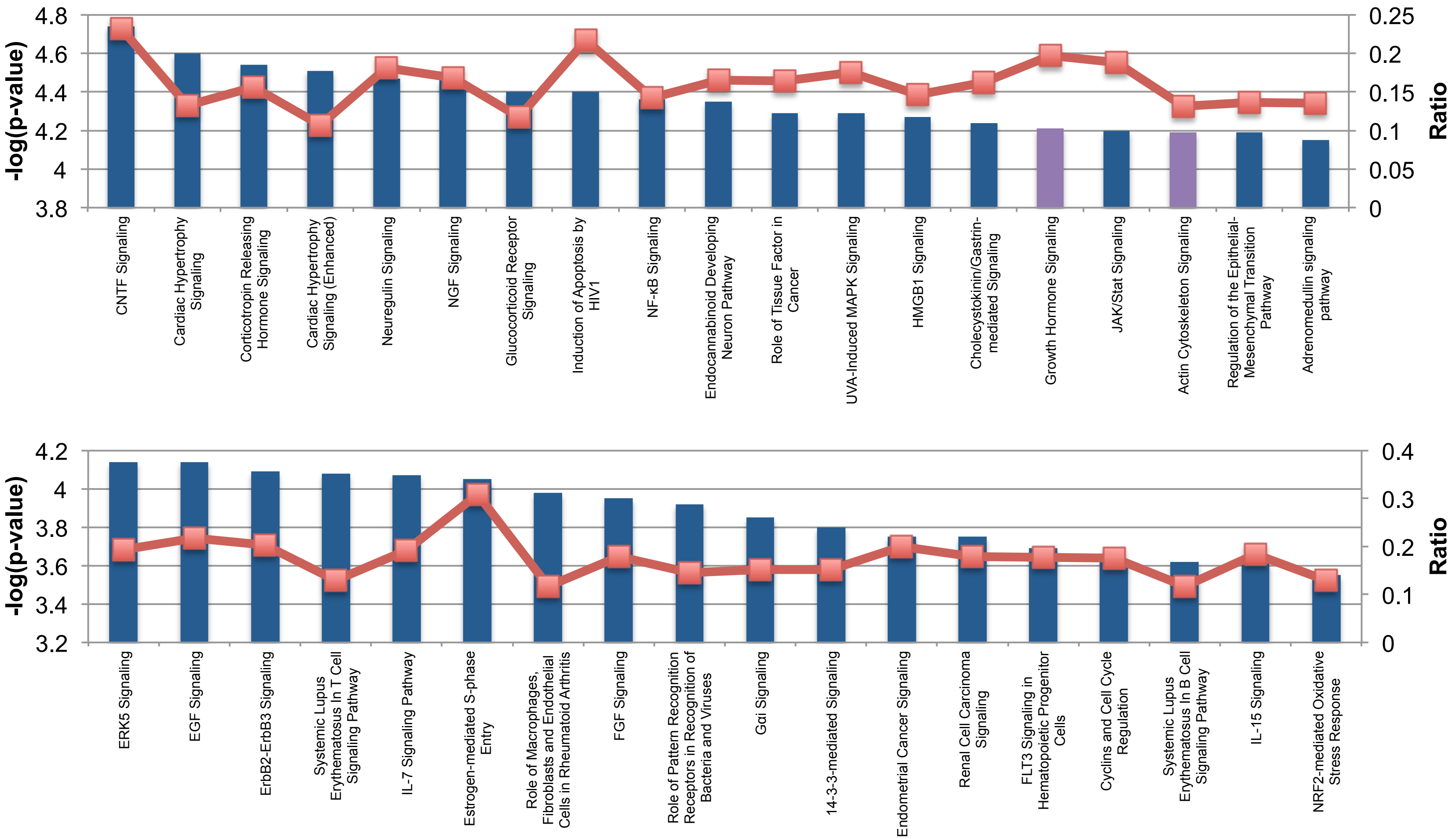
-log(p-value)
Ratio
4.8
4.6
4.4
4.2
4
3.8
0.25
0.2
0.15
0.1
0.05
0
CNTF Signaling
Cardiac Hypertrophy Signaling
Corticotropin Releasing Hormone Signaling
Cardiac Hypertrophy Signaling (Enhanced)
Neuregulin Signaling
NGF Signaling
Glucocorticoid Receptor Signaling
Induction of Apoptosis by HIV1
NF-κB Signaling
Endocannabinoid Developing Neuron Pathway
Role of Tissue Factor in Cancer
UVA-Induced MAPK Signaling
HMGB1 Signaling
Cholecystokinin/Gastrin-mediated Signaling
Growth Hormone Signaling
JAK/Stat Signaling
Actin Cytoskeleton Signaling
Regulation of the Epithelial-Mesenchymal Transition Pathway
Adrenomedullin signaling pathway
-log(p-value)
Ratio
4.2
4
3.8
3.6
3.4
3.2
0.4
0.3
0.2
0.1
0
ERK5 Signaling
EGF Signaling
ErbB2-ErbB3 Signaling
Systemic Lupus Erythematosus In T Cell Signaling Pathway
IL-7 Signaling Pathway
Estrogen-mediated S-phase Entry
Role of Macrophages, Fibroblasts and Endothelial Cells in Rheumatoid Arthritis
FGF Signaling
Role of Pattern Recognition Receptors in Recognition of Bacteria and Viruses
Gαi Signaling
14-3-3-mediated Signaling
Endometrial Cancer Signaling
Renal Cell Carcinoma Signaling
FLT3 Signaling in Hematopoietic Progenitor Cells
Cyclins and Cell Cycle Regulation
Systemic Lupus Erythematosus In B Cell Signaling Pathway
IL-15 Signaling
NRF2-mediated Oxidative Stress Response

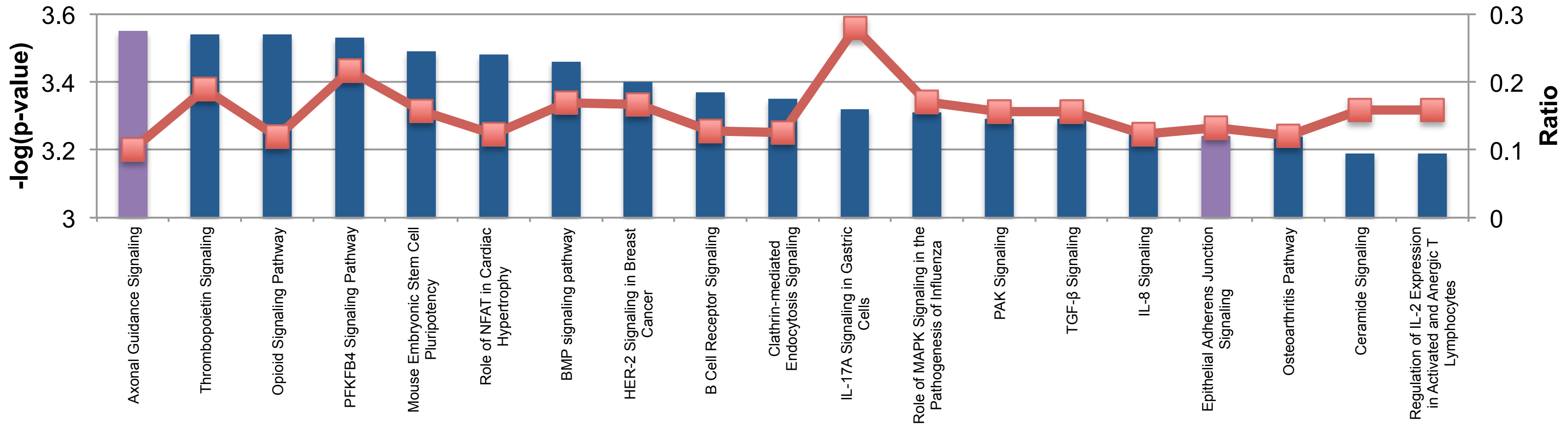

3.6
3.4
3.2
3
-log(p-value)
0.3
0.2
0.1
0
Ratio
Axonal Guidance Signaling
Thrombopoietin Signaling
Opioid Signaling Pathway
PFKFB4 Signaling Pathway
Mouse Embryonic Stem Cell Pluripotency
Role of NFAT in Cardiac Hypertrophy
BMP signaling pathway
HER-2 Signaling in Breast Cancer
B Cell Receptor Signaling
Clathrin-mediated Endocytosis Signaling
IL-17A Signaling in Gastric Cells
Role of MAPK Signaling in the Pathogenesis of Influenza
PAK Signaling
TGF-β Signaling
IL-8 Signaling
Epithelial Adherens Junction Signaling
Osteoarthritis Pathway
Ceramide Signaling
Regulation of IL-2 Expression in Activated and Anergic T Lymphocytes


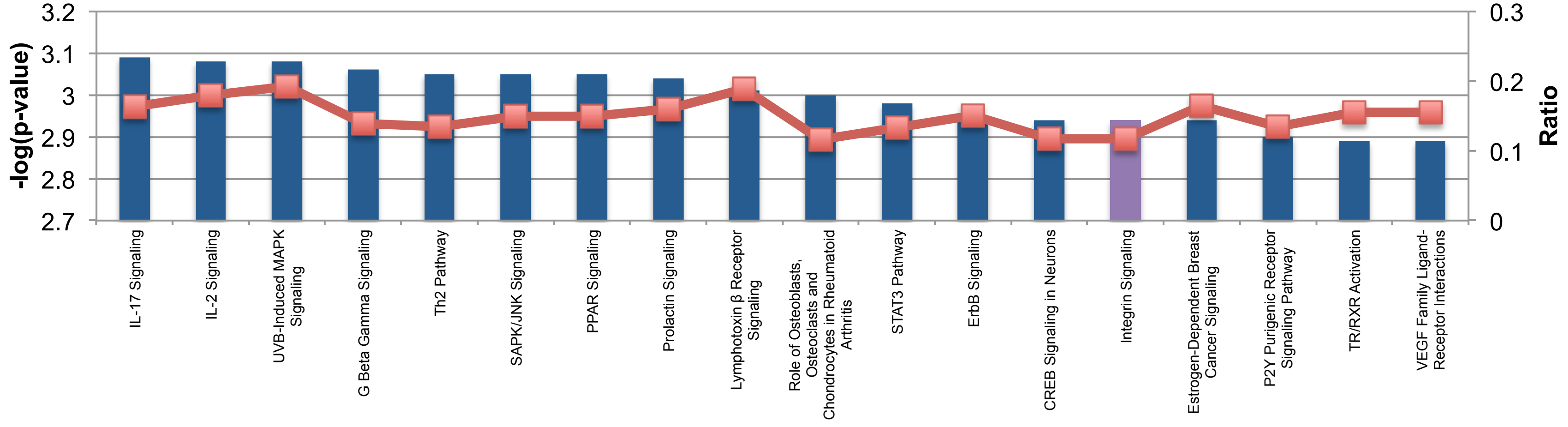

3.2
3.1
3
2.9
2.8
2.7
-log(p-value)
0.3
0.2
0.1
0
Ratio
IL-17 Signaling
IL-2 Signaling
UVB-Induced MAPK Signaling
G Beta Gamma Signaling
Th2 Pathway
SAPK/JNK Signaling
PPAR Signaling
Prolactin Signaling
Lymphotoxin β Receptor Signaling
Role of Osteoblasts, Osteoclasts and Chondrocytes in Rheumatoid Arthritis
STAT3 Pathway
ErbB Signaling
CREB Signaling in Neurons
Integrin Signaling
Estrogen-Dependent Breast Cancer Signaling
P2Y Purigenic Receptor Signaling Pathway
TR/RXR Activation
VEGF Family Ligand-Receptor Interactions

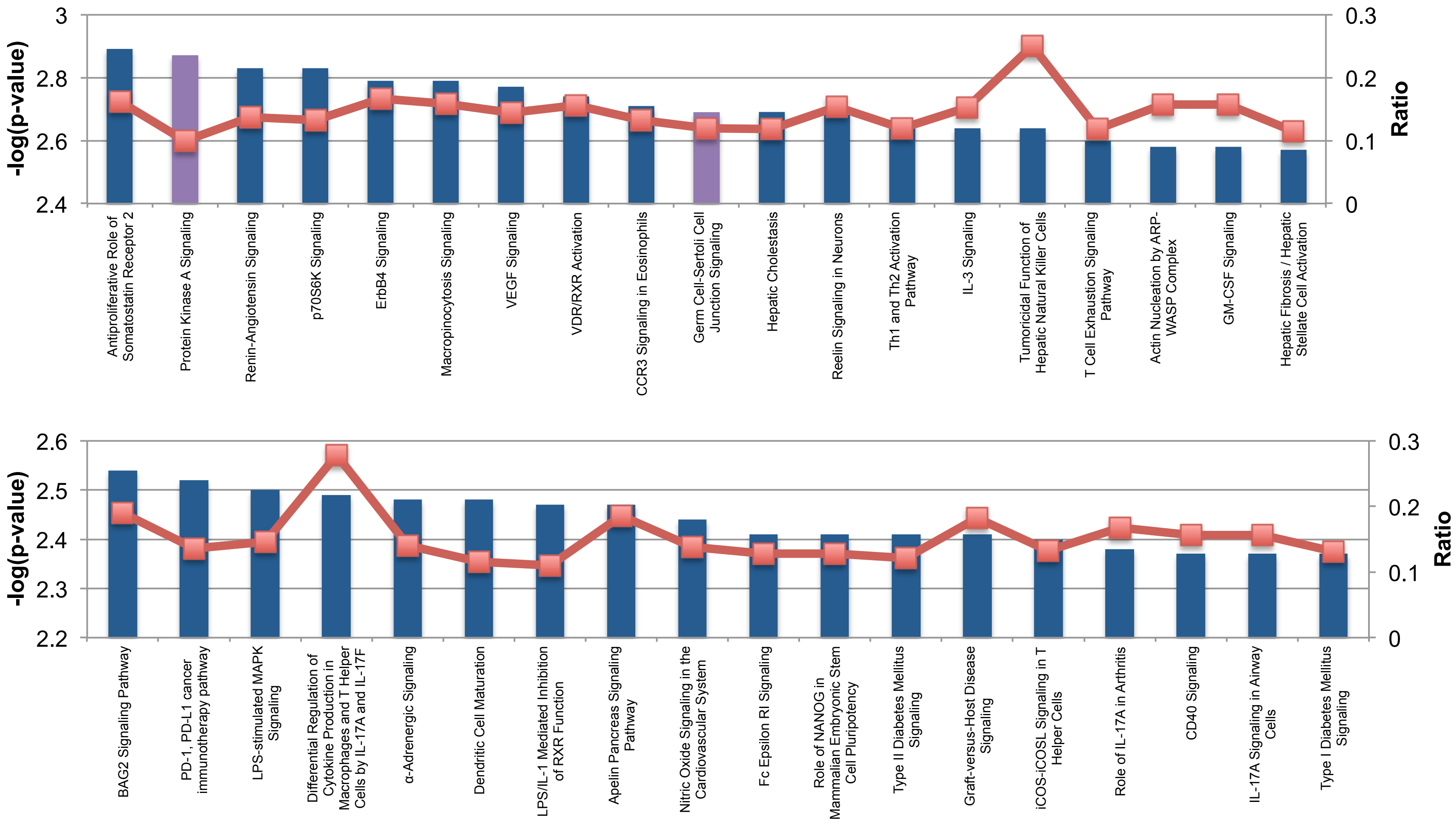
-log(p-value)
Ratio
3
2.8
2.6
2.4
0.3
0.2
0.1
0
Antiproliferative Role of Somatostatin Receptor 2
Protein Kinase A Signaling
Renin-Angiotensin Signaling
p70S6K Signaling
ErbB4 Signaling
Macropinocytosis Signaling
VEGF Signaling
VDR/RXR Activation
CCR3 Signaling in Eosinophils
Germ Cell-Sertoli Cell Junction Signaling
Hepatic Cholestasis
Reelin Signaling in Neurons
Th1 and Th2 Activation Pathway
IL-3 Signaling
Tumoricidal Function of Hepatic Natural Killer Cells
T Cell Exhaustion Signaling Pathway
Actin Nucleation by ARP-WASP Complex
GM-CSF Signaling
Hepatic Fibrosis / Hepatic Stellate Cell Activation
-log(p-value)
Ratio
2.6
2.5
2.4
2.3
2.2
0.3
0.2
0.1
0
BAG2 Signaling Pathway
PD-1, PD-L1 cancer immunotherapy pathway
LPS-stimulated MAPK Signaling
Differential Regulation of Cytokine Production in Macrophages and T Helper Cells by IL-17A and IL-17F
α-Adrenergic Signaling
Dendritic Cell Maturation
LPS/IL-1 Mediated Inhibition of RXR Function
Apelin Pancreas Signaling Pathway
Nitric Oxide Signaling in the Cardiovascular System
Fc Epsilon RI Signaling
Role of NANOG in Mammalian Embryonic Stem Cell Pluripotency
Type II Diabetes Mellitus Signaling
Graft-versus-Host Disease Signaling
iCOS-iCOSL Signaling in T Helper Cells
Role of IL-17A in Arthritis
CD40 Signaling
IL-17A Signaling in Airway Cells
Type I Diabetes Mellitus Signaling

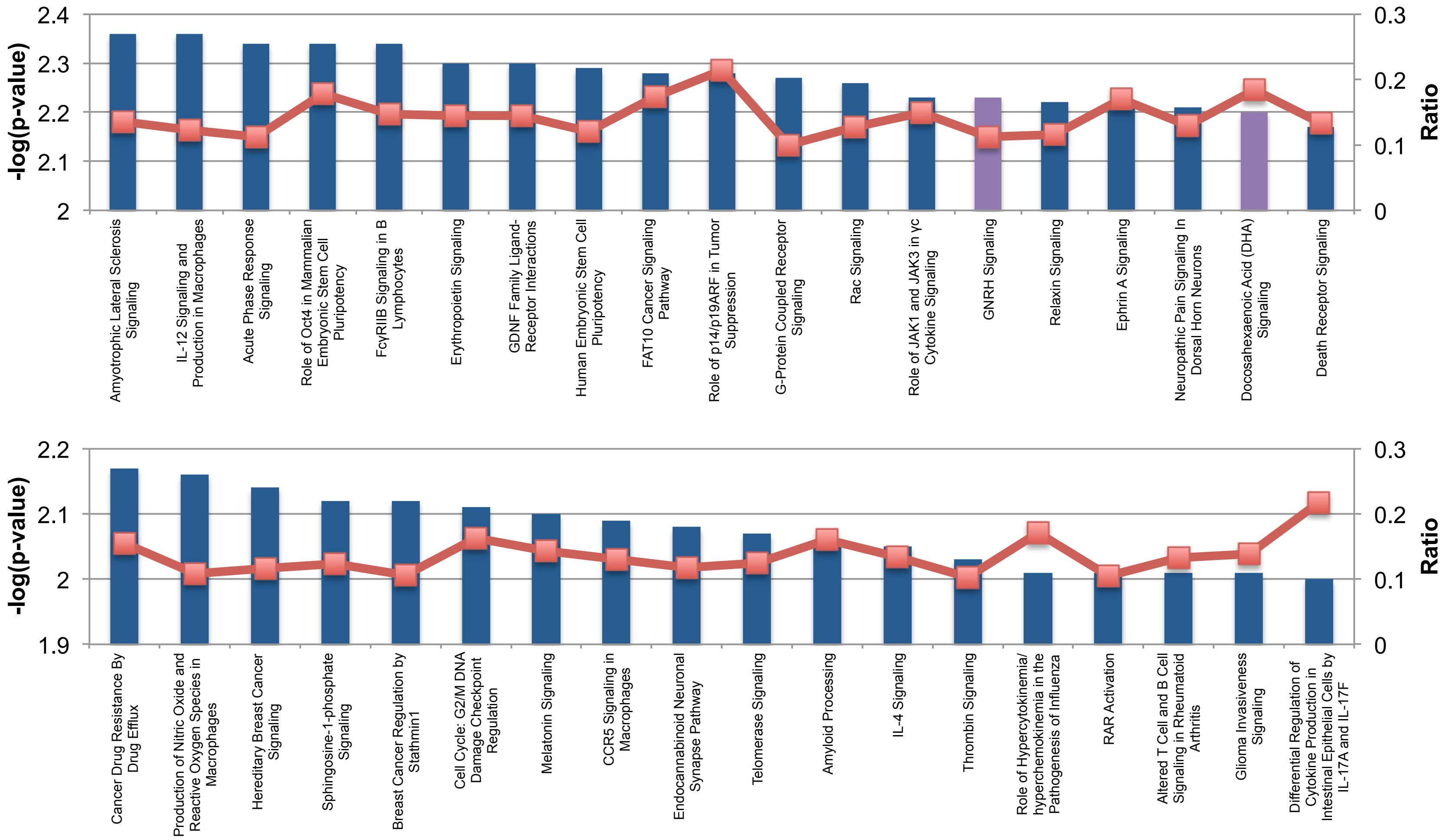

-log(p-value)
Ratio
2.4
2.3
2.2
2.1
2
0.3
0.2
0.1
0
Amyotrophic Lateral Sclerosis Signaling
IL-12 Signaling and Production in Macrophages
Acute Phase Response Signaling
Role of Oct4 in Mammalian Embryonic Stem Cell Pluripotency
FcγRIIB Signaling in B Lymphocytes
Erythropoietin Signaling
GDNF Family Ligand-Receptor Interactions
Human Embryonic Stem Cell Pluripotency
FAT10 Cancer Signaling Pathway
Role of p14/p19ARF in Tumor Suppression
G-Protein Coupled Receptor Signaling
Rac Signaling
Role of JAK1 and JAK3 in γc Cytokine Signaling
GNRH Signaling
Relaxin Signaling
Ephrin A Signaling
Neuropathic Pain Signaling In Dorsal Horn Neurons
Docosahexaenoic Acid (DHA) Signaling
Death Receptor Signaling
-log(p-value)
Ratio
2.2
2.1
2
1.9
0.3
0.2
0.1
0
Cancer Drug Resistance By Drug Efflux
Production of Nitric Oxide and Reactive Oxygen Species in Macrophages
Hereditary Breast Cancer Signaling
Sphingosine-1-phosphate Signaling
Breast Cancer Regulation by Stathmin1
Cell Cycle: G2/M DNA Damage Checkpoint Regulation
Melatonin Signaling
CCR5 Signaling in Macrophages
Endocannabinoid Neuronal Synapse Pathway
Telomerase Signaling
Amyloid Processing
IL-4 Signaling
Thrombin Signaling
Role of Hypercytokinemia/hyperchemokinemia in the Pathogenesis of Influenza
RAR Activation
Altered T Cell and B Cell Signaling in Rheumatoid Arthritis
Glioma Invasiveness Signaling
Differential Regulation of Cytokine Production in Intestinal Epithelial Cells by IL-17A and IL-17F

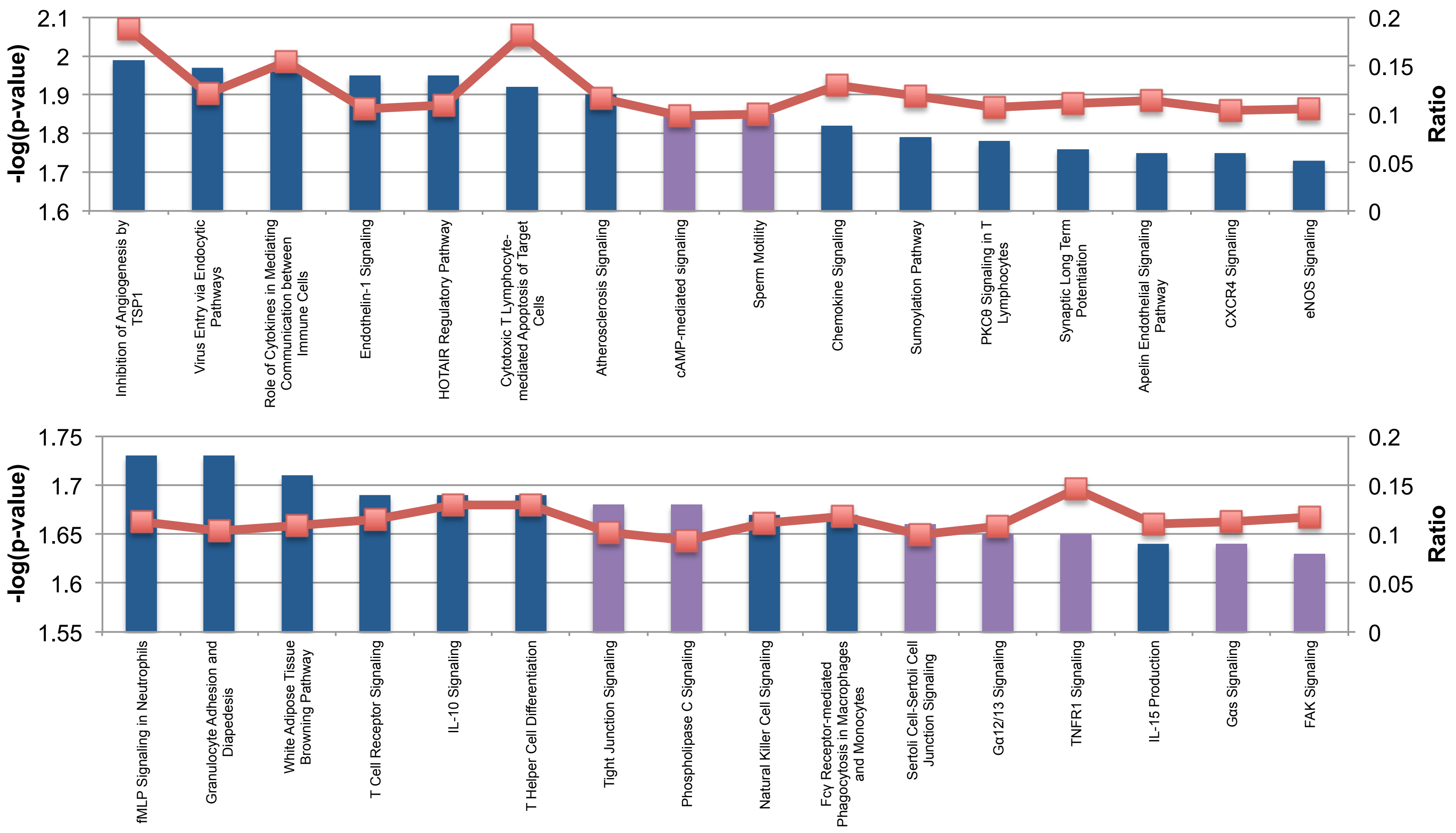

-log(p-value)
Ratio
2.1
2
1.9
1.8
1.7
1.6
0.2
0.15
0.1
0.05
0
Inhibition of Angiogenesis by TSP1
Virus Entry via Endocytic Pathways
Role of Cytokines in Mediating Communication between Immune Cells
Endothelin-1 Signaling
HOTAIR Regulatory Pathway
Cytotoxic T Lymphocyte-mediated Apoptosis of Target Cells
Atherosclerosis Signaling
cAMP-mediated signaling
Sperm Motility
Chemokine Signaling
Sumoylation Pathway
PKCθ Signaling in T Lymphocytes
Synaptic Long Term Potentiation
Apelin Endothelial Signaling Pathway
CXCR4 Signaling
eNOS Signaling
-log(p-value)
Ratio
1.75
1.7
1.65
1.6
1.55
0.2
0.15
0.1
0.05
0
fMLP Signaling in Neutrophils
Granulocyte Adhesion and Diapedesis
White Adipose Tissue Browning Pathway
T Cell Receptor Signaling
IL-10 Signaling
T Helper Cell Differentiation
Tight Junction Signaling
Phospholipase C Signaling
Natural Killer Cell Signaling
Fcγ Receptor-mediated Phagocytosis in Macrophages and Monocytes
Sertoli Cell-Sertoli Cell Junction Signaling
Gα12/13 Signaling
TNFR1 Signaling
IL-15 Production
Gαs Signaling
FAK Signaling

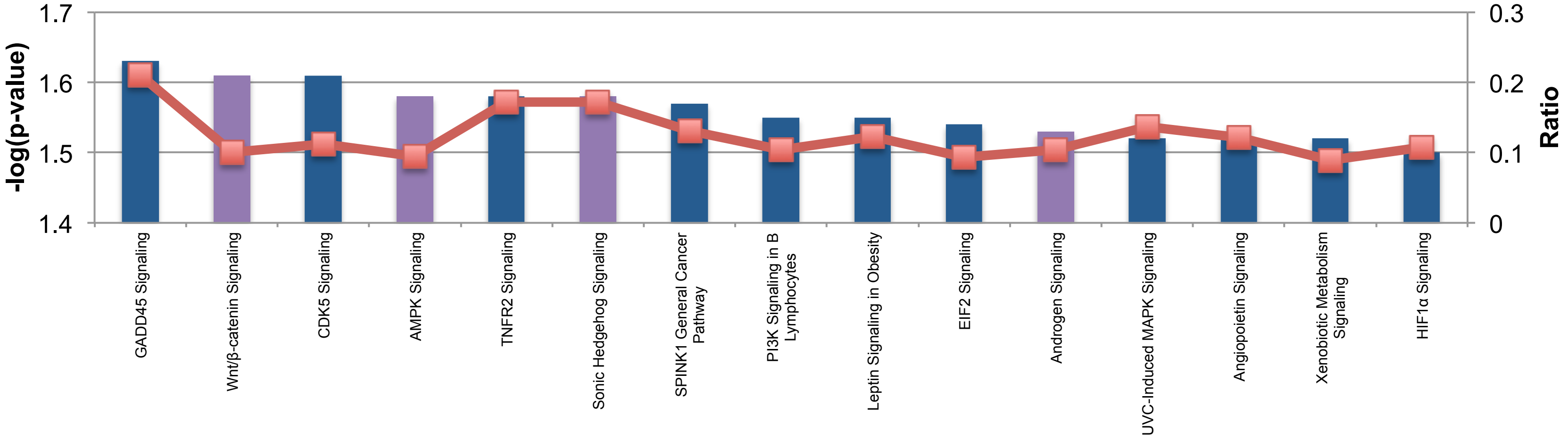


Highlighted in purple are the known sperm concentration related gene pathways. The dotted red line represents the ratio of the miRNA target genes to the total number of genes known to be present in that pathway.

**Supplemental Table S1: Top 10 pathways (FDR q-value < 0.05) enriched in 234 genes associated with baseline piRNA listed in Supplemental File S5, sheet 6**

| Gene Set Name | Description | # Genes in Overlap (k) | k/K | p-value | FDR q-value |
|---|---|---|---|---|---|
| GO_GOLGI_CIS_CISTERNA | The Golgi cisterna closest to the endoplasmic reticulum; the first processing compartment through which proteins pass after export from the ER. [ISBN:0815316194] | 15 | 0.5357 | 1.84E-27 | 2.29E-23 |
| GO_CIS_GOLGI_NETWORK | The network of interconnected tubular and cisternal structures located at the convex side of the Golgi apparatus, which abuts the endoplasmic reticulum. [ISBN:0198506732, ISBN:0815316194] | 15 | 0.2206 | 1.8E-20 | 1.13E-16 |
| GO_GOLGI_CISTERNA_MEMBRANE | The lipid bilayer surrounding any of the thin, flattened compartments that form the central portion of the Golgi complex. [GOC:ecd, GOC:mah] | 16 | 0.1739 | 5.73E-20 | 2.38E-16 |
| GO_GOLGI_CISTERNA | Any of the thin, flattened membrane-bounded compartments that form the central portion of the Golgi complex. [GOC:mah] | 16 | 0.1379 | 2.82E-18 | 8.8E-15 |
| GO_GOLGI_ORGANIZATION | A process that is carried out at the cellular level which results in the assembly, arrangement of constituent parts, or disassembly of the Golgi apparatus. [GOC:dph, GOC:jl, GOC:mah] | 16 | 0.1127 | 7.81E-17 | 1.95E-13 |
| GO_GOLGI_STACK | The set of thin, flattened membrane-bounded compartments, called cisternae, that form the central portion of the Golgi complex. The stack usually comprises cis, medial, and trans cisternae; the cis- and trans-Golgi networks are not considered part of the stack. [GOC:mah, ISBN:0815316194] | 16 | 0.1067 | 1.9E-16 | 3.95E-13 |
| GO_ACTIVATION_OF_GTPASE_ACTIVITY | Any process that initiates the activity of an inactive GTPase through the replacement of GDP by GTP. [GOC:dph, GOC:mah, GOC:tb] | 11 | 0.1146 | 4.79E-12 | 8.53E-09 |
| GO_RAB_GTPASE_BINDING | Interacting selectively and non-covalently with Rab protein, any member of the Rab subfamily of the Ras superfamily of monomeric GTPases. [GOC:mah] | 12 | 0.0682 | 2.39E-10 | 3.73E-07 |
| GO_ENDOMEMBRANE_SYSTEM_ORGANIZATION | A process that is carried out at the cellular level which results in the assembly, arrangement of constituent parts, or disassembly of the endomembrane system. [GOC:mah, GOC:sm] | 17 | 0.039 | 2.76E-10 | 3.83E-07 |
| GO_ORGANELLE_SUBCOMPARTMENT | A compartment that consists of a lumen and an enclosing membrane, and is part of an organelle. [GOC:mah, GOC:pz] | 16 | 0.0419 | 3.35E-10 | 4.17E-07 |

**Supplemental Table S2. Baseline tRFs and piRNAs associated with BMI and sperm concentration at -log10 adjusted p-value ≥ 1.3**

**A** tRFs associated with BMI

| tRFs | General tRF name | logFC | AveExpr | t | P.Value | adj.P.Val | B |
|---|---|---|---|---|---|---|---|
| chr17-tir5-33-GlnTTG | tir5-GlnTTG | -0.593 | 11.541 | -5.005 | 0.000 | 0.004 | 2.465 |
| chr17-trf5c-33-GlnTTG | trf5c-GlnTTG | -0.542 | 13.024 | -4.470 | 0.000 | 0.009 | 1.127 |
| chr17-trf5b-33-GlnTTG | trf5b-GlnTTG | -0.589 | 10.939 | -4.165 | 0.000 | 0.014 | 0.376 |
| chr1-tir5-127-CysGCA | tir5-CysGCA | -0.312 | 5.605 | -3.852 | 0.001 | 0.023 | -0.356 |
| chr6-trf5c-84-GlnTTG | trf5c-GlnTTG | -0.307 | 5.733 | -3.637 | 0.001 | 0.033 | -0.871 |
| chr7-trf5c-7-CysGCA | trf5c-CysGCA | -0.273 | 5.585 | -3.473 | 0.002 | 0.042 | -1.255 |

**B** tRFs associated with sperm concentration

| tRFs | General tRF name | logFC | AveExpr | t | P.Value | adj.P.Val | B |
|---|---|---|---|---|---|---|---|
| chr6-trf5c-70-AlaCGC | trf5c-AlaCGC | -0.036 | 11.602 | -5.005 | 0.000 | 0.005 | 1.861 |
| chr6-trf5b-110-AlaTGC | trf5b-AlaTGC | -0.043 | 4.015 | -4.475 | 0.000 | 0.009 | 0.607 |
| chr6-tir5-87-GluCTC | tir5-GluCTC | -0.028 | 15.802 | -4.200 | 0.000 | 0.013 | -0.349 |
| chr6-tir5-152-ValCAC | tir5-ValCAC | 0.027 | 9.524 | 3.894 | 0.001 | 0.018 | -1.018 |
| chr2-trf5c-27-GlyCCC | trf5c-GlyCCC | 0.029 | 6.627 | 3.867 | 0.001 | 0.018 | -1.051 |
| chr19-trf5c-2-GlyTCC | trf5c-GlyTCC | 0.023 | 9.050 | 3.444 | 0.002 | 0.046 | -2.127 |

**C** piRNAs associated with sperm concentration

| piRNAs | logFC | AveExpr | t | P.Value | adj.P.Val | B |
|---|---|---|---|---|---|---|
| piR_002703 | -0.031 | 6.694 | -5.847 | 0.000 | 0.008 | 3.901 |
| piR_018904 | -0.027 | 6.980 | -5.558 | 0.000 | 0.009 | 3.133 |
| piR_001455 | 0.026 | 7.139 | 4.938 | 0.000 | 0.029 | 1.425 |

**Supplemental Table S3. Diet intervention induced alterations in sncRNA expression**

**A**

| miRNAs | logFC | AveExpr | t | P.Value | adj.P.Val | B |
|---|---|---|---|---|---|---|
| miR-513c-3p | 2.535 | 1.021 | 3.823 | 0.000 | 0.501 | -1.295 |
| miR-4774-3p | -6.385 | 2.180 | -3.459 | 0.001 | 0.508 | -1.848 |
| miR-509-3-5p | 1.647 | 9.282 | 3.426 | 0.001 | 0.508 | -1.204 |
| miR-1229-3p | -4.566 | 1.471 | -3.213 | 0.003 | 0.508 | -2.362 |
| miR-656-5p | -3.741 | 1.029 | -3.164 | 0.003 | 0.508 | -2.382 |
| miR-4760-5p | -1.911 | 0.561 | -3.152 | 0.003 | 0.508 | -2.389 |
| miR-506-3p | 2.404 | 6.121 | 3.147 | 0.003 | 0.508 | -1.992 |
| miR-503-5p | 2.337 | 2.740 | 3.068 | 0.004 | 0.551 | -2.582 |
| miR-6511b-5p | -2.106 | 1.034 | -2.979 | 0.005 | 0.599 | -2.704 |
| miR-513a-3p | 2.082 | 0.970 | 2.955 | 0.005 | 0.599 | -2.662 |
| miR-136-3p | 2.046 | 3.294 | 2.894 | 0.006 | 0.642 | -2.722 |
| miR-508-3p | 1.634 | 7.588 | 2.834 | 0.007 | 0.651 | -2.460 |
| miR-6503-3p | -4.031 | 2.466 | -2.825 | 0.007 | 0.651 | -2.769 |
| miR-4507 | -1.881 | 5.591 | -2.747 | 0.009 | 0.718 | -2.730 |
| miR-3147 | -3.035 | 1.538 | -2.701 | 0.010 | 0.718 | -3.066 |

**B**

| tRFs | General tRF name | logFC | AveExpr | t | P.Value | adj.P.Val | B |
|---|---|---|---|---|---|---|---|
| chr2-trf5b-2-TyrGTA | trf5b-TyrGTA | 3.231 | 1.356 | 3.302 | 0.002 | 0.437 | -4.557 |
| chr17-tir5-29-CysGCA | tir5-CysGCA | -1.477 | 9.047 | -2.851 | 0.007 | 0.437 | -4.491 |
| chr6-trf5b-24-AlaAGC | trf5b-AlaAGC | -2.297 | 4.641 | -2.850 | 0.007 | 0.437 | -4.541 |

A) The differentially expressed miRNAs (intervention vs. control) B) The differentially expressed tRFs (intervention vs. control). These DE sncRNAs were identified after adjusting for age, BMI, sperm concentration and sperm motility using log2(1.5) fold change cut-off and an un-adjusted -log10 p-value ≥2

#### Supplemental Table S4. Pathway analysis of miRNA targeted genes altered by diet intervention

**A**

| KEGG pathway | p-value | #genes | #miRNAs |
|---|---|---|---|
| Fatty acid biosynthesis | 3E-14 | 5 | 5 |
| Prion diseases | 1E-09 | 10 | 8 |
| Mucin type O-Glycan biosynthesis | 3E-06 | 13 | 7 |
| Signaling pathways regulating pluripotency of stem cells | 3E-06 | 60 | 12 |
| Proteoglycans in cancer | 3E-06 | 76 | 13 |
| TGF-beta signaling pathway | 6E-06 | 37 | 11 |
| Fatty acid metabolism | 0.0001 | 15 | 7 |
| Biotin metabolism | 0.0005 | 2 | 4 |
| Hippo signaling pathway | 0.0007 | 60 | 14 |
| Renal cell carcinoma | 0.0016 | 29 | 11 |
| Long-term depression | 0.0031 | 27 | 13 |
| Wnt signaling pathway | 0.0044 | 49 | 10 |
| FoxO signaling pathway | 0.0055 | 54 | 15 |
| Prolactin signaling pathway | 0.0114 | 29 | 10 |
| Adherens junction | 0.0298 | 31 | 11 |
| Focal adhesion | 0.0298 | 74 | 12 |
| Insulin signaling pathway | 0.0319 | 53 | 10 |
| Transcriptional misregulation in cancer | 0.0319 | 58 | 13 |
| Oxytocin signaling pathway | 0.0319 | 54 | 13 |
| Oocyte meiosis | 0.0415 | 40 | 12 |

(A) Gene pathways identified using DIANA mirPath software, at a FDR corrected p-value of <0.05.
(B) mirPath micro-T CDS target prediction scores for 7 DE-miRNAs targeting fatty acid metabolism genes.

Note: Highlighted in purple are the miRNAs known to be involved in fatty acid biosynthesis and metabolism. There is an overlap in between the miRNAs in the two highlighted pathways; a total of 7 unique miRNAs. The targets of the 7 miRNAs are show in the Table B.

**B**

**miRNAs and target genes involved in Fatty Acid Metabolism**

**microT-CDS Predicted Interactions for miR-136-3p**

| # | Gene Name | Gene Ensembl id | Score |
|---|---|---|---|
| 1 | SCD | ENSG00000099194 | 0.824 |

**microT-CDS Predicted Interactions for miR-506-3p**

| # | Gene Name | Gene Ensembl id | Score |
|---|---|---|---|
| 1 | ACAA2 | ENSG00000167315 | 0.948 |
| 2 | ACSL1 | ENSG00000151726 | 0.853 |
| 3 | CPT1A | ENSG00000110090 | 0.994 |
| 4 | ELOVL5 | ENSG00000012660 | 0.981 |
| 5 | HADH | ENSG00000138796 | 0.822 |
| 6 | OXSM | ENSG00000151093 | 0.856 |
| 7 | PECR | ENSG00000115425 | 0.901 |
| 8 | SCD | ENSG00000099194 | 0.925 |

**microT-CDS Predicted Interactions for miR-508-3p**

| # | Gene Name | Gene Ensembl id | Score |
|---|---|---|---|
| 1 | ACOX1 | ENSG00000161533 | 0.894 |

**microT-CDS Predicted Interactions for miR-509-3-5p**

| # | Gene Name | Gene Ensembl id | Score |
|---|---|---|---|
| 1 | ACADVL | ENSG00000072778 | 0.983 |
| 2 | ACSL6 | ENSG00000164398 | 0.951 |
| 3 | PTPLB | ENSG00000206527 | 0.917 |

**microT-CDS Predicted Interactions for miR-513a-3p**

| # | Gene Name | Gene Ensembl id | Score |
|---|---|---|---|
| 1 | ACADSB | ENSG00000196177 | 0.834 |
| 2 | ACOX1 | ENSG00000161533 | 0.814 |
| 3 | ELOVL5 | ENSG00000012660 | 0.834 |
| 4 | FASN | ENSG00000169710 | 0.829 |
| 5 | PTPLB | ENSG00000206527 | 0.952 |

**microT-CDS Predicted Interactions for miR-513c-3p**

| # | Gene Name | Gene Ensembl id | Score |
|---|---|---|---|
| 1 | ACADSB | ENSG00000196177 | 0.837 |
| 2 | ACOX1 | ENSG00000161533 | 0.814 |
| 3 | ELOVL5 | ENSG00000012660 | 0.832 |
| 4 | FASN | ENSG00000169710 | 0.831 |
| 5 | PTPLB | ENSG00000206527 | 0.952 |

**microT-CDS Predicted Interactions for miR-4760-5p**

| # | Gene Name | Gene Ensembl id | Score |
|---|---|---|---|
| 1 | ACSL3 | ENSG00000123983 | 0.818 |
| 2 | ELOVL5 | ENSG00000012660 | 0.879 |

**List of Supplemental Files**

**Supplemental File S1. Subject characteristics and demographics, sperm analysis, semen analysis and blood measures of the study participants**

Sheet 1: sncRNA sub set (N=17) pre-intervention

Sheet 2: Full set (N=102) pre-intervention

Sheet 3: sncRNA sub set (N=17) post-intervention

Sheet 4: Full set (N=102) post-intervention

**Supplemental File S2. Pre-intervention baseline characterization of sncRNA landscape**

Sheet 1: Expression profile of 401 mature miRNAs

Sheet 2: Expression profile of 143 tRFs

Sheet 3: Expression profile of 2290 piRNAs

**Supplemental File S3. Baseline tRF sequences present in the pre-intervention samples**

Sheet 1: Most abundant sequence for each tRF ID across the 17 samples.

Sheet 2: Representative tRF sequence for each tRF ID

**Supplemental File S4. Homer annotation of all the genomic locations of baseline piRNAs**

Sheet 1: Baseline piRNAs overlapping with transposable elements (TEs)

Sheet 2: Baseline piRNAs overlapping with tRNAs

Sheet 3: Baseline piRNAs not overlapping with transposable elements (NTEs)

**Supplemental File S5. Mapping of piRNABank identified baseline piRNA to piRBase annotations**

Sheet 1: piRNABank piRNA identifiers that mapped to piRBase

Sheet 2: LINE related baseline piRNAs in piRBase

Sheet 3: SINE related baseline piRNAs in piRBase

Sheet 4: LTR related baseline piRNAs in piRBase

Sheet 5: Satellite related baseline piRNAs in piRBase

Sheet 6: Genes related baseline piRNAs in piRBase

**Supplemental File S6. miRNA associated with age and sperm concentration**

Sheet 1: miRNAs associated with age (adjusted –log10 p value ≥ 1.3)

Sheet 2: miRNAs associated with sperm concentration (adjusted –log10 p value ≥ 1.3)

**Supplemental File S7. Differentially expressed sncRNA between control and intervention**

sncRNA highlighted in purple represent the DE-sncRNA identified after adjusting for the effects of age, BMI, sperm concentration and motility.

Sheet 1: Differentially expressed miRNA (unadjusted –log10 p value ≥ 2).

Sheet 2: Differentially expressed tRF (unadjusted –log10 p value ≥ 2).

Sheet 3: Differentially expressed piRNA (unadjusted –log10 p value ≥ 2).

**Supplemental File S8: Homer annotation of all the genomic locations of DE-piRNAs**

Sheet 1: DE-piRNAs overlapping with transposable elements (TEs)

Sheet 2: DE-piRNAs not overlapping with transposable elements (NTEs)

**Supplemental File S9: Mapping of piRNABank identified DE-piRNA to piRBase annotations**

Sheet 1: piRNABank piRNA identifiers that mapped with piRBase.

Sheet 2: Repeats related DE-piRNAs in piRBase

Sheet 3: Genes related DE-piRNAs in piRBase
